# HIV-1-induced cytokines deplete homeostatic ILCs and expand TCF7-dependent memory NK cells

**DOI:** 10.1101/221010

**Authors:** Yetao Wang, Kyle Gellatly, Sean McCauley, Pranitha Vangala, Kyusik Kim, Alan Derr, Smita Jaiswal, Alper Kucukural, Patrick McDonel, Lawrence Lifshitz, Thomas Greenough, JeanMarie Houghton, Manuel Garber, Jeremy Luban

## Abstract

HIV-1-infected people who take medications that suppress viremia, preserve CD4^+^ T cells, and prevent AIDS, have chronic inflammation with increased cardiovascular mortality. To investigate the etiology of this inflammation, the effect of HIV-1 on innate lymphoid cells (ILCs) and NK cells was examined. Homeostatic ILCs in blood and intestine were depleted permanently. NK cells were skewed towards a memory subset. Cytokines that are elevated during HIV-1 infection reproduced both abnormalities *ex vivo*. Pseudotime analysis of single NK cell transcriptomes revealed a developmental trajectory towards a subset with expression profile, chromatin state, and biological function like memory T lymphocytes. Expression of TCF7, a WNT transcription factor, increased over the course of the trajectory. TCF7 disruption, or WNT inhibition, prevented memory NK cell induction by inflammatory cytokines. These results demonstrate that inflammatory cytokines associated with HIV-1 infection irreversibly disrupt homeostatic ILCs and cause developmental shift towards TCF7^+^ memory NK cells.

**Highlights:**

- HIV-1 infection depletes homeostatic ILCs in blood and intestine and shifts NK cells towards a memory cell phenotype, irrespective of viremia or CD4 count
- Inflammatory cytokines recapitulate ILC and NK cell abnormalities ex vivo
- TCF7 expression correlates with a developmental trajectory that culminates in memory NK cells
- TCF7/WNT signaling is required for establishment of memory NK cells

Graphical Abstract

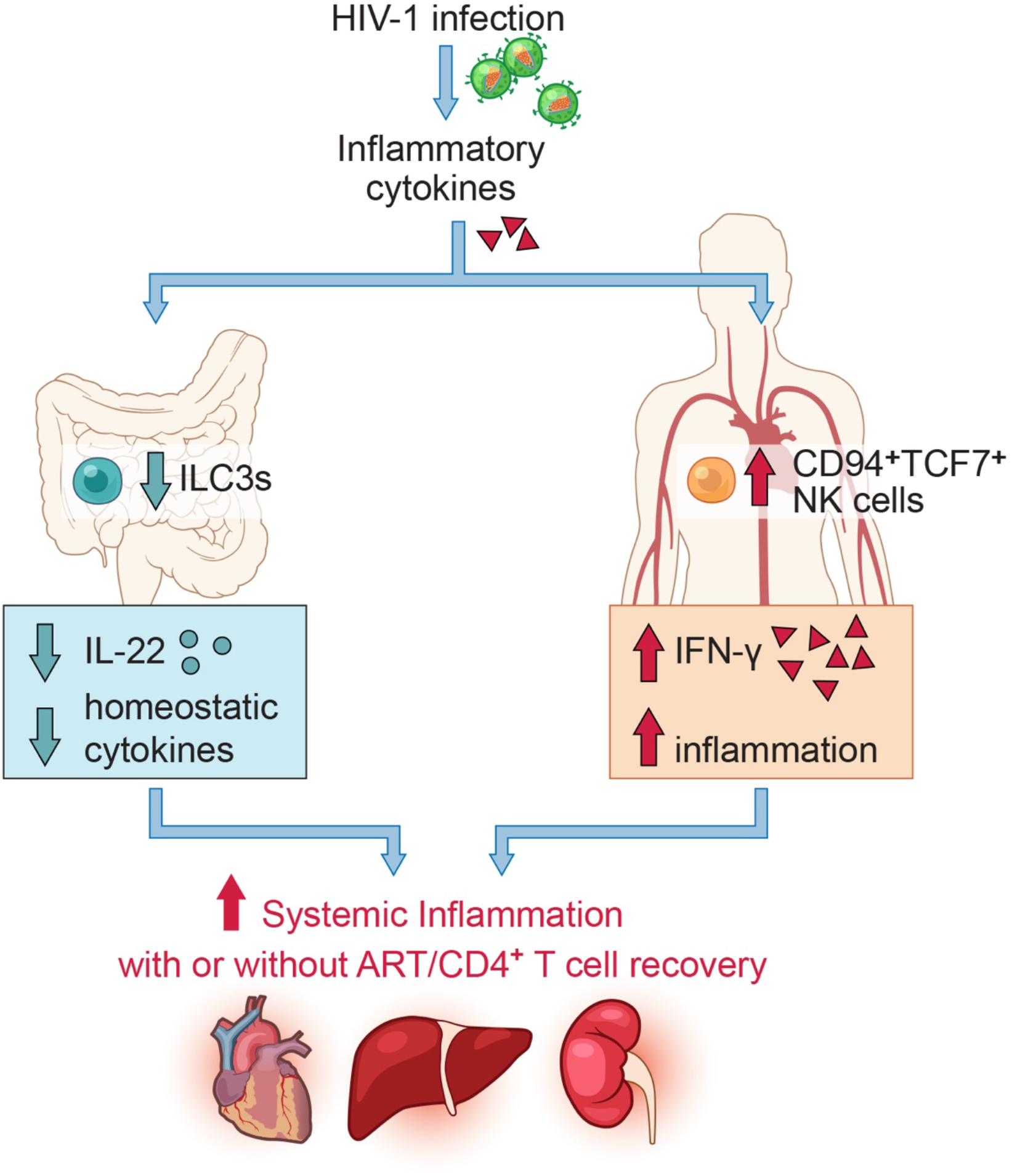

## INTRODUCTION

HIV-1 infection is lifelong and no treatments yet available reliably eliminate the virus from infected people (Margolis et al., 2016). Given uninterrupted access to antiviral drug combinations that suppress HIV-1 viremia it is possible to preserve CD4^+^ T cells and prevent progression to AIDS (Günthard et al., 2016). Nonetheless, infected individuals on these antiviral therapies have chronic inflammation with increased rates of cardiovascular mortality (Freiberg et al., 2013; Sinha et al., 2016). This pathology has been attributed to permanent disruption of cellular networks that maintain the integrity of the intestinal epithelium and limit the intestinal microbiome (Brenchley et al., 2006; Deeks et al., 2013a; Klatt et al., 2012).

Innate lymphoid cells (ILCs) are counterparts of T cells that lack clonotypic antigen receptors or other lineage-defining cell surface markers (Hazenberg and Spits, 2014; Spits et al., 2013). ILCs carry out a large range of biological functions, that include roles in host defense against pathogens and homeostatic maintenance in inflamed tissues (Artis and Spits, 2015; Klose and Artis, 2016). Among the many subsets of ILCs are natural killer (NK) cells that are capable of lysing tumor cells and virus-infected cells (Spits et al., 2016; Vivier et al., 2008). ILCs are divided into three main subtypes based on key transcription factors and the cytokines that they produce. ILC1s and NK cells both express TBX21 and secrete IFN-γ. ILC2s require GATA3 for their development and produce IL-4, IL-5, IL-9, IL-13 and Amphiregulin. ILC3s express RORγT and produce IL-17 and IL-22 (Spits et al., 2013; Walker et al., 2013).

Blood ILCs are irreversibly depleted by HIV-1 if antiviral therapy is not started within a few weeks of acute infection (Kløverpris et al., 2016). The contribution of ILC deficiency to HIV-1-associated chronic inflammation remains unclear since abnormalities in homeostatic ILCs from tissues such as the colon lamina propria have not been reported. NK cells kill HIV-1-infected cells and can eliminate infected cells via antibody-dependent cell cytotoxicity (Alter et al., 2011; Bruel et al., 2016; Haynes et al., 2012). The clinical significance of NK cells for control of HIV-1 is supported by the fact that genetic association between HLA haplotypes on target cells and KIRs on NK cells contribute to viral control during HIV-1 infection (Lin et al., 2016b; Martin et al., 2002; Pelak et al., 2011).

NK cells have historically been recognized as innate immune counterparts of cytotoxic CD8^+^ T cells that rapidly respond to infected cells or tumors, independent of antigen-specific recognition. Studies in mice, rhesus macaques, and humans, though, have shown that the response of NK cells can be increased by prior encounter with inflammatory cytokines, haptens, vaccination, or viruses (Cerwenka and Lanier, 2016; O’Sullivan et al., 2015). Whether these memory-like NK cell populations share transcriptional and chromatin features with memory T cells has not been thoroughly investigated.

In this study, the effect of chronic HIV-1 infection on ILCs and NK cells was examined. HIV-1 infection was found to decrease the numbers of all ILCs in human blood and in colon lamina propria, and to alter the balance between CD94^-^ and CD94^+^ NK cells. Deeper investigation revealed that TCF7^+^CD94^+^CD56^hi^ NK cells are distinct from other NK populations in that they exhibit functional, transcriptional, and epigenetic features characteristic of CD8^+^ memory T cells, and that such memory NK cells are expanded by HIV-1 infection.

## RESULTS

### HIV-1 Infection Decreases Homeostatic ILCs

To identify ILCs from among human peripheral blood mononuclear cells (PBMCs), 12 cell surface proteins were used to exclude T cells, B cells, NK cells and other cells from defined lineages within the lymphoid gate (Figure S1A) (Hazenberg and Spits, 2014; Kløverpris et al., 2016; Lim et al., 2017). Then, CD127^+^ cells from the Lin^-^ population were assessed for TBX21 to distinguish ILC1s, CRTH2 for ILC2s, and RORγT for ILC3s (Figure S1B) (Hazenberg and Spits, 2014). Lin^-^CD127^+^ ILC1s and ILC3s, capable of producing IFN-γ or IL-22, respectively, were rare in the blood (Figure S1B). The majority of the Lin^-^CD127^+^ cells in HIV-1^-^ blood were CRTH2^+^ ILC2s, capable of producing IL-13 (Figure S1B) (Colonna, 2018; Lim et al., 2017), and all Lin^-^CD127^+^ILCs were decreased in the blood of people with HIV-1 infection (Figures 1A and 1B; Table S1). ILC reduction was significant (p<0.01) whether all HIV-1^+^ samples were grouped together (Figure 1B), or samples were stratified based on viremia or CD4 count (Figures S1C and S1D).

**Figure 1.**
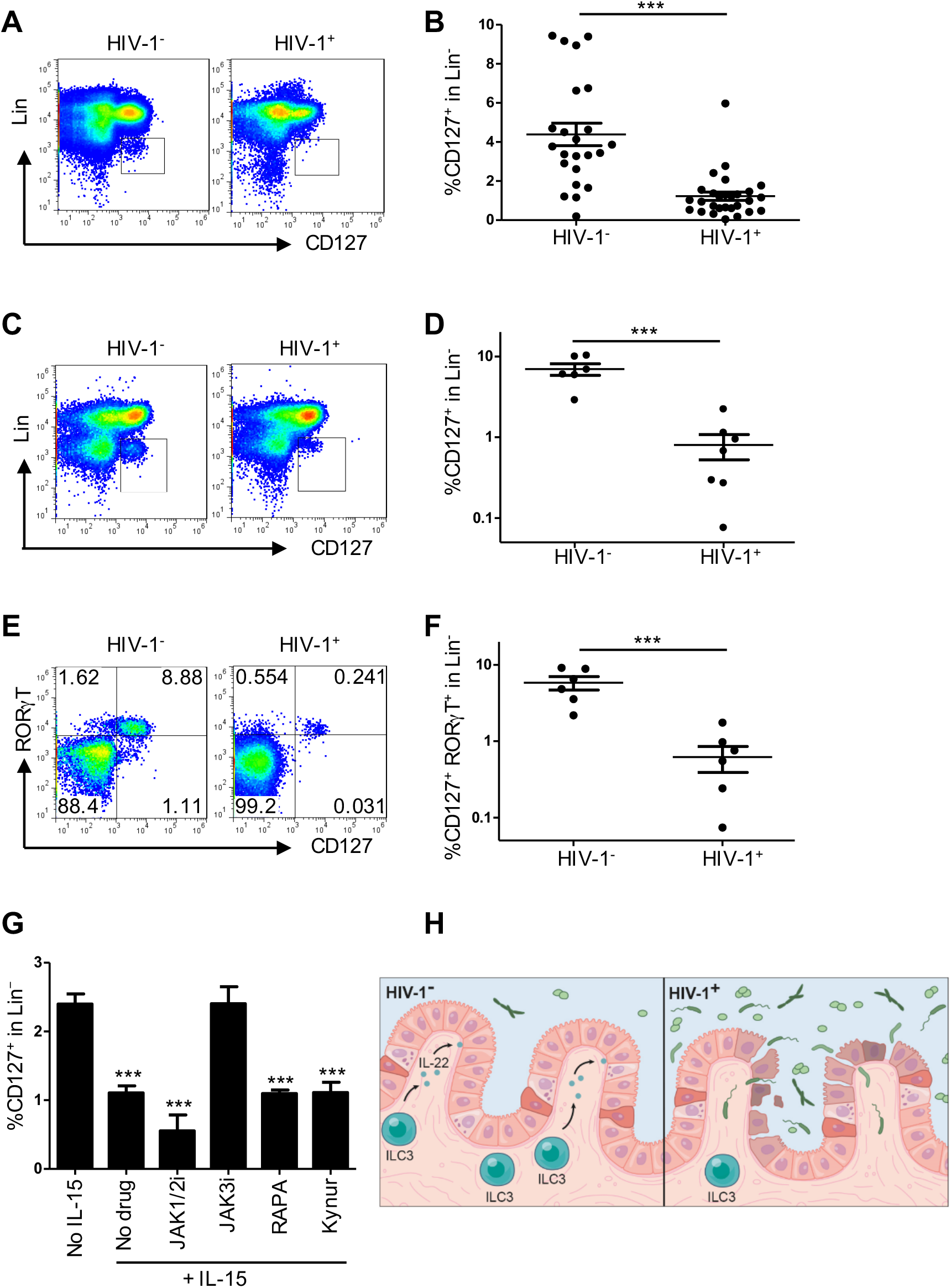
HIV-1 Infection Decreases ILCs in Blood and Colon Lamina Propria. (A) Flow cytometry for 12 lineage markers versus CD127 on live, singlet, lymphoid PBMCs from HIV-ľ and HIV-1^+^ individuals. (B) Percent ILCs (Lin^-^CD127^+^ PBMCs) from HIV-ľ (n=23) and HIV-1^+^ (n=28) individuals, gated as in (A). (C) Colon lamina propria cells, gated as in (A). (D) As in (C), percent ILCs (Lin^-^CD127^+^ colon lamina propria lymphoid cells) from HIV-ľ (n=6) and HIV-1^+^ individuals on antiretroviral therapy with undetectable viremia (n=7). (E) Lin^-^ colon lamina propria cells showing CD127 versus RORγT. (F) Percent ILC3s (Lin^-^CD127^+^RORyT^+^ colon lamina propria cells) from HIV-ľ (n=6) and HIV-1^+^ (n=7) individuals, as in (E). (G) Percent CD127^+^ cells among Lin^-^ PBMCs from HIV-ľ donors (n=4) after incubation with IL-15 for 16 hrs in the presence of ruxolitinib (JAK1/2i), CP-690550 (JAK3i), rapamycin (RAPA), or L-kynurinine (Kynur). (H) IL-22 produced by lamina propria ILC3s maintains gut epithelium integrity (left). Irreversible decrease in ILC3s with HIV-1 infection (right). Data is mean ± s.e.m., two-tailed unpaired ř-test; ***p<0.001. See also Figure S1.

Chronic systemic inflammation in HIV-1^+^ people has been associated with inappropriate translocation of bacterial products from the intestinal lumen (Brenchley et al., 2006; Deeks et al., 2013a). Given the reduced number of ILCs in blood from HIV-1 ^+^ individuals, and the fact that lamina propria ILCs contribute to the maintenance of gut epithelial integrity (Artis and Spits, 2015), the effect of HIV-1 infection on colon lamina propria ILCs was examined next. In HIV-1^-^ controls, the majority of Lin^-^CD127^+^ colon ILCs were RORγT^+^ and capable of producing IL-22 in response to stimulation (Figures 1C-1F and S1E). In HIV-1^+^ individuals on anti-HIV-1 therapy, colon lamina propria ILC3s were reduced in number (Figures 1C-1F), despite undetectable viremia in these individuals, and no significant reduction in lamina propria CD4^+^ T cells (Figure S1F; Table S1).

ILCs do not express CD4 or CCR5, they do not bear these proteins on their surface, and they cannot be infected *ex vivo* by HIV-1 (Figure S1G) (Kløverpris et al., 2016). Reduction in ILCs from HIV-1^+^ people is therefore unlikely to result from direct infection of these cells by HIV-1. To determine whether the decrease in CD127^+^ILCs associated with HIV-1 infection might be a consequence of systemic elevation in cytokines, inflammatory metabolites, or leakage of microbes across the intestinal epithelium (Brenchley et al., 2006; Deeks et al., 2013a; Favre et al., 2010; Kacani et al., 1997; Roberts et al., 2010; Shebl et al., 2012; Swaminathan et al., 2016), the effect on Lin^-^CD127^+^PBMCs of exposure to these factors *in vitro* was assessed. No significant decrease in CD127 was observed after exposure to any of 13 cytokines, chemokines, TLR agonists, bacterial products, or L-kynurinine (Figures S1H-S1J). However, as reported for T lymphocytes (Mazzucchelli and Durum, 2007), common Y-chain cytokines that are systemically elevated during HIV-1 infection (Kacani et al., 1997; Roberts et al., 2010; Shebl et al., 2012; Stacey et al., 2009; Swaminathan et al., 2016), including IL-2, IL-4, and IL-15, downregulated CD127 on ILCs (Figures S1K). A JAK3 inhibitor, CP-690550, prevented CD127 downregulation by IL-15 (Figure 1G), consistent with a requirement for cytokine signaling via JAK3 (Leonard and O’Shea, 1998). Neither the JAK1/2-inhibitor ruxolitinib nor the MTOR-inhibitor rapamycin prevented CD127 downregulation (Figure 1G). These results suggest that JAK3 signaling, in response to systemic elevation of common γ-chain cytokines, decreases CD127^+^ILCs in HIV-1 infection. This would deprive intestinal epithelium of homeostatic ILC3s, disrupt the integrity of the colon epithelium, and explain ongoing inflammation associated with HIV-1 infection (Figure 1H).

### HIV-1 Infection Increases the Proportion of CD94^+^ NK Cells

CD94 is found on a subset of NK cells (Chang et al., 1995). When the anti-CD94 antibody was removed from the panel of lineage antibodies used above, all Lin^-^TBX21 ^+^ PBMCs, whether CD94^+^ or CD94^-^, were CD56^+^EOMES^+^NK cells (Figures S2A and S2B). When HIV-1^-^ and HIV-1^+^ individuals were compared for these markers, no significant difference in the percentage of total Lin^-^TBX21^+^NK cells was detected (Figures 2A and 2B). In contrast, the ratio of CD94^+^/CD94^-^NK cells was significantly increased in PBMCs from HIV-1^+^ people (Figures 2C and 2D). Consistent with this observation, when CD94^+^ cells were excluded with the other lineage markers, Lin^-^ TBX21^+^ cells were decreased in HIV-1^+^ people (Figure S2C). These findings regarding the effect of HIV-1 on NK cells were independent of the magnitude of viremia or the number of blood CD4^+^ T cells (Figures S2D and S2E).

**Figure 2.**
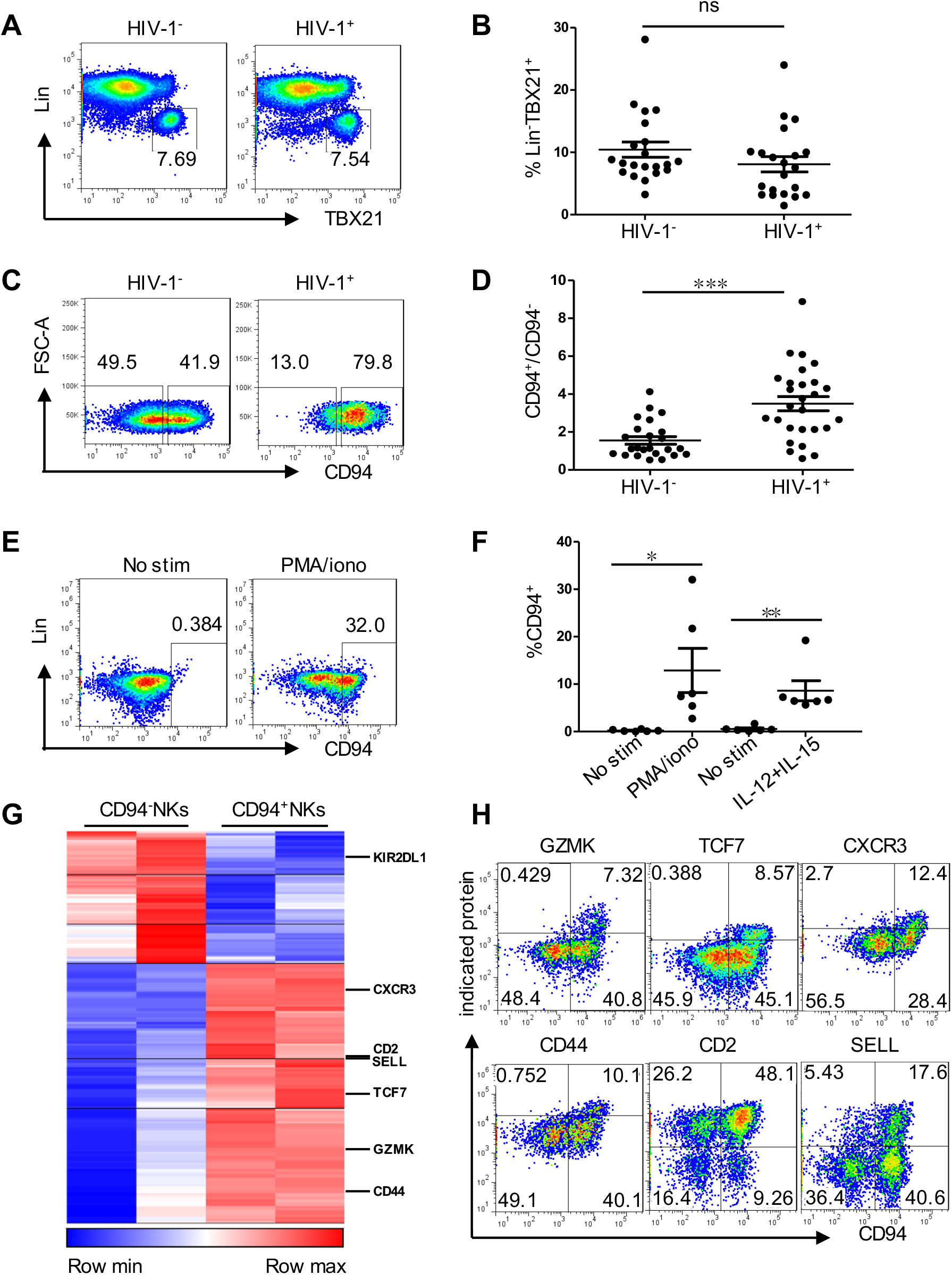
HIV-1 Infection Increases the Proportion of CD94^+^ NK Cells. (A) NK cells (Lin^-^TBX21^+^ PBMCs) from HIV-1^-^ and HIV-1^+^ individuals. (B) As in (A), percent NK cells from HIV-1^-^(n=21) and HIV-1^+^ (n=21) individuals. (C) CD94 on Lin^-^TBX21^+^ PBMCs from HIV-1^-^ and HIV-1^+^ individuals. (D) As in (C), fraction of CD94^+^NK cells among Lin^-^TBX21^+^ PBMCs from HIV-1^-^ (n=24) and HIV-1^+^ (n=27) individuals. (E) Percent CD94^+^ cells after 16 hrs PMA/ionomycin stimulation of Lin^-^CD56+CD94^-^NK cells sorted from HIV-1^-^ PBMCs. (F) As in (E), percent CD94^+^ NK cells after 16 hrs stimulation with PMA/ionomycin or IL-12 and IL15, n=6 HIV-1^-^ donors. (G) Heatmap of genes that are differentially expressed in Lin^-^CD56+CD94^-^ NK versus Lin^-^CD56+CD94^+^NK cells, sorted from PBMCs (two donors), as determined by RNA-Seq (fold change, log2 >1, p<0.05). (H) Flow cytometry for proteins encoded by differentially expressed genes as in (G). Data is mean ± s.e.m. (B), two-tailed unpaired *t*-test; (D), Mann-Whitney test; (F), two-tailed paired *t*-test; ns, not significant; *p<0.05, **p<0.01, ***p<0.001. See also Figure S2.

Stimulation of PBMCs with PMA and ionomycin, with IL-15 alone, or with IL-15 plus IL-12, increased the ratio of CD94^+^ to CD94^-^NK cells, similar to that observed in the blood of HIV-1^+^ people (Figure S2F). When CD94^-^NK cells were sorted (Lin^-^CD56^+^CD94^-^PBMCs in Figure S2G), CD94 levels and degranulation activity increased in response to stimulation with PMA and ionomycin, or IL-12 and IL-15 (Figures 2E, 2F, and S2H). In the absence of stimulation, sorted CD94^+^NK cells had greater cytolytic activity than did CD94^-^NK cells (Figure S2I). Levels of Ki-67 and Annexin V were comparable on the CD94^-^ and CD94^+^NK cells, suggesting that the perturbed ratio of these cells *in vivo* did not result from intrinsic differences in rates of proliferation or apoptosis (Figures S2J).

### TCF7 Expression Tracks with Pseudotime Trajectory from CD94^-^CD56^dim^NK Cells to CD94^+^CD56^hi^NK Cells

To better understand the molecular basis for the increase in CD94^+^NK cells associated with HIV-1 infection, transcriptional profiles of sorted CD94^-^ and CD94^+^ NK cells were obtained (Figure S2G). RNA-Seq revealed 140 genes expressed at higher level in CD94^+^ NK cells (Figure 2G; Table S2) and proteins encoded by differentially-expressed genes were confirmed by flow cytometry (Figures 2H and S2K). CD94^+^NK cells expressed higher levels of GZMK (p=5.64E-04), TCF7 (p=1.68E-11), CXCR3 (p=2.51-07), CD44 (p=6.45E-12), CD2 (p=4.59E-06) and SELL (p=1.63E-05), the biological functions of which include NK cell survival (Jeevan-Raj et al., 2017), activation (Sconocchia et al., 1997; Wendel et al., 2008) and memory (Juelke et al., 2010; Liu et al., 2016). In contrast, expression (p=2.75E-05) and surface protein levels of KIR2DL1, an HLA-Cw4 ligand that inhibits NK cell cytotoxicity (Gazit et al., 2004), were increased on CD94^-^NK cells (Figures 2G and S2K).

Transcriptomes across the spectrum from CD94^-^NK to CD94^+^NK cells were then captured by single-cell RNA-Seq (scRNA-Seq) (Figures 3A and S3A; Table S3) (Klein et al., 2015). Based on the analysis of the 3,277 single cell transcriptomes, CD94^-^NK cells formed a homogeneous population (Figure 3A, pink dots in upper right). CD94^+^NK cells formed two populations, one of which overlapped with CD94^-^NK cells (Figure 3A, aqua dots, upper right). Unbiased clustering of total Lin^-^CD56^+^ PBMCs, irrespective of whether a cell was CD94^+^, also showed two distinct cell populations (Figure 3B). The validity of clustering cells into two groups was confirmed by calculating prediction strength as a function of cluster number (Tibshirani and Walther, 2005) (Figure S3B). Cluster 1 was homogeneous and consisted of 396 CD94^+^NK cells (Figure 3B, red dots lower left). Cluster 2 contained 371 CD94^+^NK cells and 973 CD94^-^NK cells (Figure 3B, blue dots, upper right). A heatmap based on all differentially expressed genes between the two clusters showed a shift in the pattern of gene expression along the continuum of CD94 expression (see blue and yellow bars at the top of Figure 3C; Table S4).

**Figure 3.**
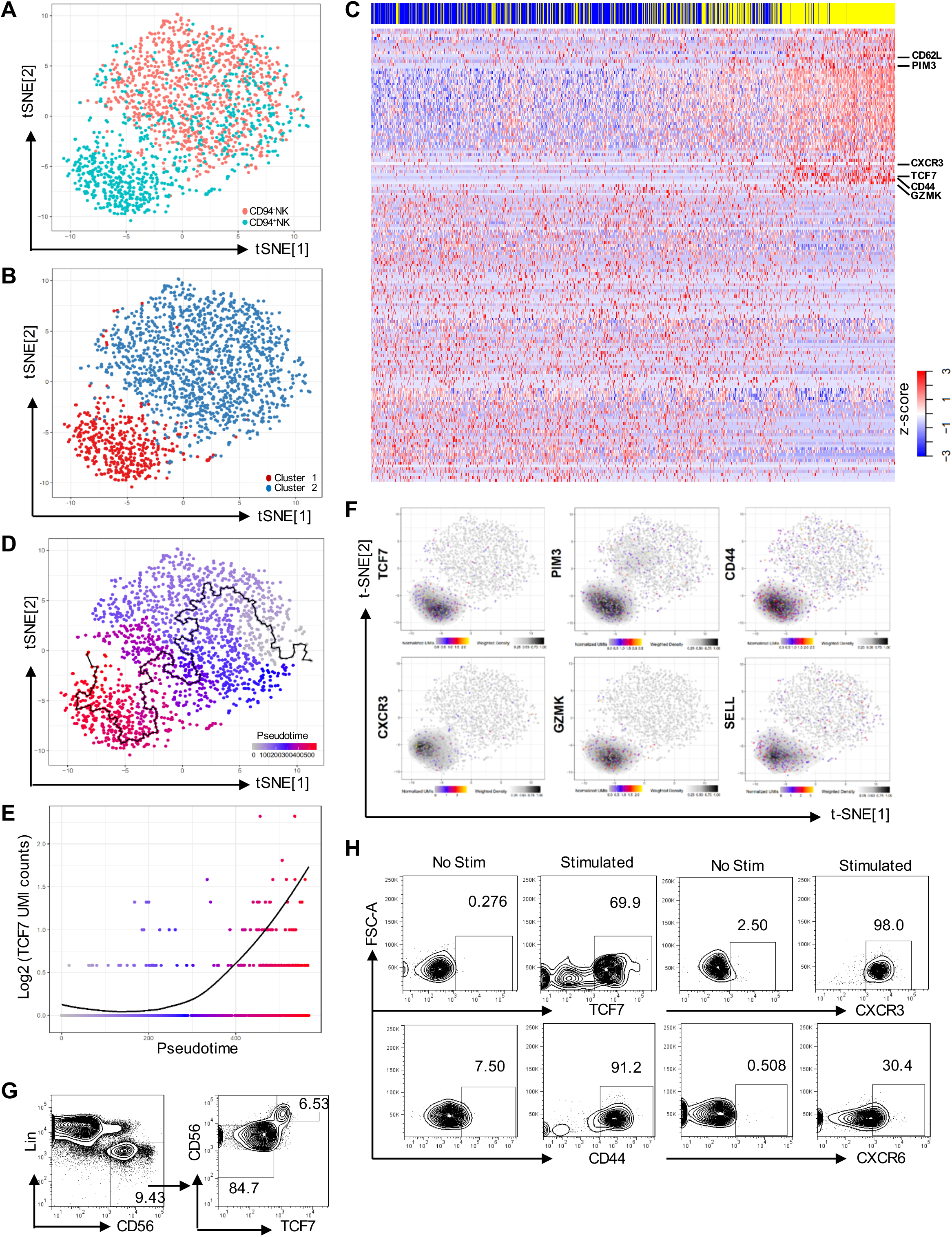
TCF7 Expression Correlates with Pseudotime Trajectory from CD94^-^NK Cells to CD94^+^NK Cells. (A) Two-dimensional tSNE plot of single cell RNA-Seq of 986 Lin^-^CD56^+^CD94^-^NK cells (pink) and 767 Lin^-^CD56^+^CD94^+^NK cells (aqua). (B) Spectral clustering of single cell transcriptomes from all cells in (A), independent of CD94, k-nearest neighbor, search=2. (C) Heatmap of 1,729 Lin^-^CD56^+^CD94^-^NK cells (blue) and 1,548 Lin^-^CD56^+^CD94^+^NK cells (yellow), collected from 2 donors, using all differentially expressed genes from the spectral cluster analysis. (D) Minimum spanning tree based on the transcriptome of individual cells from (A), showing pseudotime trajectory (black line). (E) TCF7 expression along the pseudotime trajectory. (F) Expression and density of the indicated genes within t-SNE plots. (G) Flow cytometry for CD56 and TCF7 on Lin^-^PBMCs. (H) Sorted Lin^-^CD56^+^CD94^-^NK cells were untreated (No Stim) or treated with IL-15 (5 ng/ml) for 5 days, and IL-12 (50 ng/ml) and IL-15 (50 ng/ml) for 16 hrs (Stimulated). TCF7, CD44, CXCR3, and CXCR6 were detected by flow cytometry. TCF7, representative of 8 donors; CD44, CXCR3, and CXCR6, representative of 4 donors. See also Figure S3.

To determine whether heterogeneity in the individual CD94^+^NK cell transcriptomes reflected different stages in the CD94^-^ to CD94^+^NK cell transition, a minimum spanning tree based on individual cell transcriptomes was constructed using Monocle (Trapnell et al., 2014) (Figure 3D). A heatmap utilizing the pseudotime ordering of single cells, based on the minimum spanning tree (Figure S3C), mimicked the heatmap ordered by hierarchical clustering (Figure 3C), validating the ordering of cells along the transcriptional timeline assigned by the minimum spanning tree.

Genes with potential to regulate the transition from CD94^-^NK to CD94^+^ NK cells were identified by differential expression analysis based on the pseudotemporal ordering of cells from the minimum spanning tree. Candidate genes included TCF7 (Figures 3E and 3F, false discovery rate p < 7 x 10^-103^), a transcription factor activated by WNT-signaling that is important for T cell memory establishment and maintenance (Jeannet et al., 2010; Utzschneider et al., 2016), PIM3, a serine-threonine kinase that blocks apoptosis and promotes self-renewal (Aksoy et al., 2007), CD44, a gene important for T cell survival and establishment of memory cells (Baaten et al., 2010), CXCR3 and GZMK, genes highly expressed in memory CD8^+^ T cells (Weng et al., 2012), and SELL, a marker for NK cells with potential to proliferate and differentiate into effectors upon secondary stimulation (Juelke et al., 2010). TCF7^+^NK cells examined by flow cytometry were exclusively CD56^hi^ (Figure 3G), a marker for NK cells with capacity to proliferate and differentiate into CD56^dim^ NK cells (Moretta, 2010). CXCR6, a gene required for memory NK cell generation in response to haptens and viruses (Paust et al., 2010) was also highly enriched on TCF7^+^NK cells, as were CD44, SELL, and CXCR3 (Figure S3D).

When total PBMC or sorted CD94^-^NK cells were treated with IL-15, CD94 was upregulated within 16 hrs. After 5 days of IL-15 treatment, TCF7, CXCR3, CD44, CXCR6, and GZMK were also upregulated (Figures 3H and S3E). Additionally, as previously reported (Freud et al., 2014), high levels of CD56 were observed on the cell surface (Figures S3F and S3G). These experiments stimulating cells directly *ex vivo* substantiate the pseudotemporal analysis of scRNA-Seq data, demonstrate that CD94^+^CD56^hi^ NK cells are generated from CD94^-^NK cells in response to inflammatory cytokines, and suggest that the expanded CD94^+^NK cell population in blood from HIV-1^+^ people results from elevated inflammatory cytokines.

### Distinct Chromatin Landscape of CD94^+^CD56^hi^NK Cells

To further investigate the TCF7^+^CD56^hi^ subset of CD94^+^ NK cells identified above (Figures 3B, 3F, and 3G), CD94^-^CD56^dim^, CD94^+^CD56^dim^, and CD94^+^CD56^hi^ NK cells were sorted (Figure 4A) and subjected to RNA-Seq. The transcriptional profile of CD94^+^CD56^hi^ NK cells was distinct from that of either CD94^-^CD56^dim^ or CD94^+^CD56^dim^ cells (Figure 4B), with 275 (Figure 2C; Table S5), and 162 (Figure 2D; Table S5), differentially expressed genes, respectively. In contrast, only 4 genes distinguished CD94^+^CD56^dim^ from CD94^-^CD56^dim^ NK cells (Figure 4E; Table S5). Principal component analysis (PCA) isolated the CD94^+^CD56^hi^ NK cell transcriptome as a distinct cluster (Figure 4F) and reactome pathway analysis showed enrichment for WNT signaling and TCF7 (Figure 4G and Table S5).

**Figure 4.**
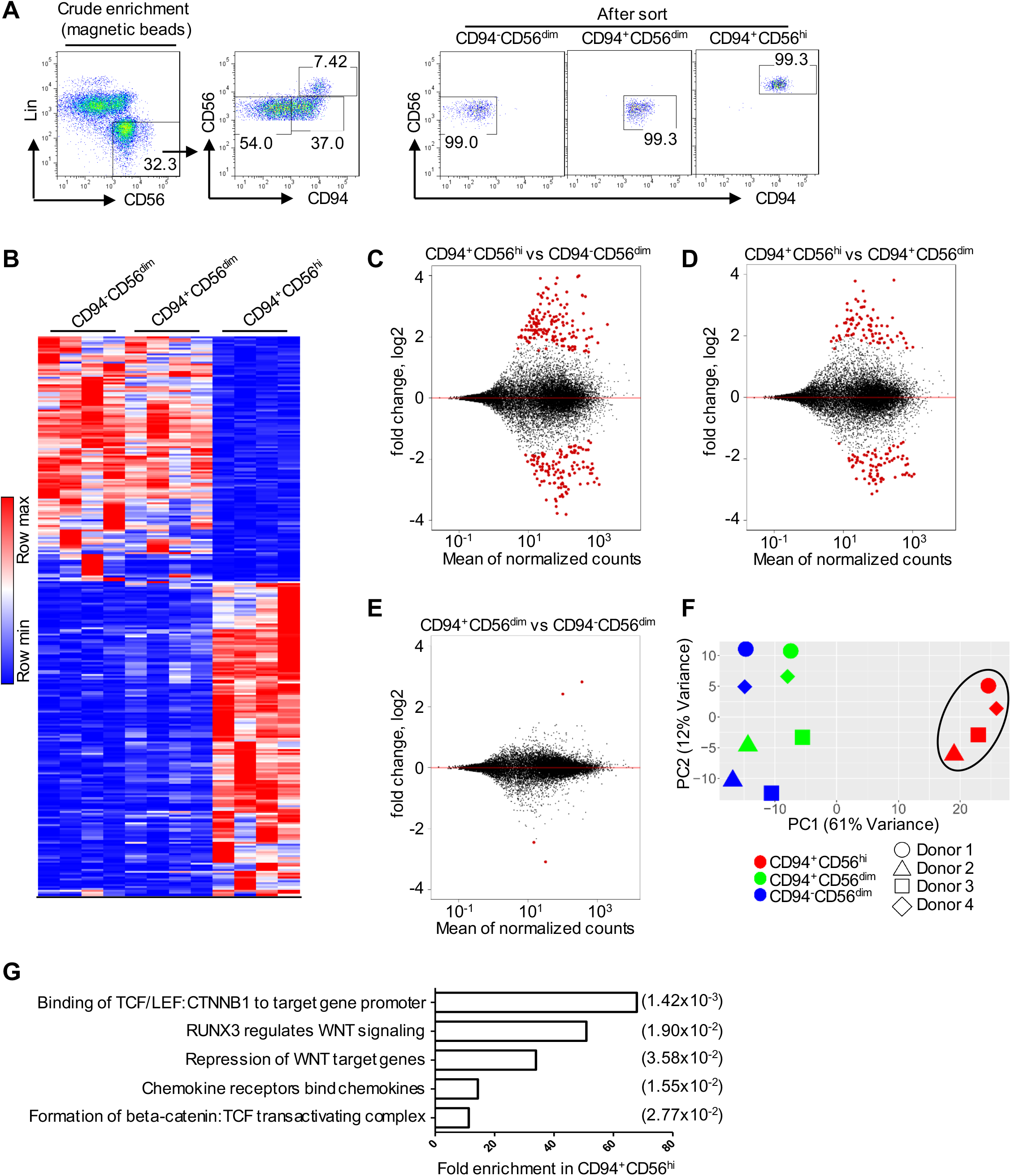
TCF/WNT Signaling Pathway Enrichment in CD94^+^CD56^hi^NK Cells. (A) Sorting strategy for CD94^-^CD56^dim^, CD94^+^CD56^dim^, and CD94^+^CD56^hi^ NK cell subsets. (B) Heatmap of differentially expressed genes by RNA-Seq (fold change of normalized counts, log2 >1, p<0.05) for the indicated NK cell subsets sorted from four PBMC donors. (C-E) Pairwise comparison of the indicated NK cell subsets based on differentially expressed genes. (F) PCA based on RNA-Seq data from the indicated NK cell subsets. (G) Reactome pathway analysis based on 152 genes that are differentially expressed in CD94^+^CD56^hi^ NK cells, with false discovery rate in parentheses.

In addition to having a unique transcriptional profile, activating histone marks and accessible chromatin regions in CD94^+^CD56^hi^ NK cells localized to chromosomal locations that were distinct from those in the other two NK subsets (Figures 5A-C, S4A, and S4B). The chromatin landscape for representative genes, including TCF7, a gene uniquely expressed in the CD94^+^CD56^hi^ NK subset, and for CD6, a gene only expressed in CD56^dim^ NK cells, are shown (Figure 5D, E and S4C). *De novo* motif analysis showed enrichment for TCF7, RUNX, NK-κB, and four other DNA binding motifs within the accessible chromatin regions that were unique to CD94^+^CD56^hi^ NK cells (Figure 5F). In contrast, the IKZF1 binding motif was enriched within both CD56^dim^ NK cell subsets (Figure 5G). TCF7 binding peaks mapped to 255 CD94^+^CD56^hi^ NK cell ATAC-Seq peaks located within genes encoding cytokine receptors IL2RA, IL2RB, and IL20RA, transcription factors RUNX1, RUNX3, and NOTCH2, chromatin modifiers SETD5 and SUV420H2 (KMT5C), and the WNT signaling regulator AXIN1 (Figures 5H and S4D). These transcriptional and epigenetic profiles indicate that, among NK cells, CD94^+^CD56^hi^ NK cells constitute a developmentally discrete subset, and that TCF7 is critical for the establishment of this population.

**Figure 5.**
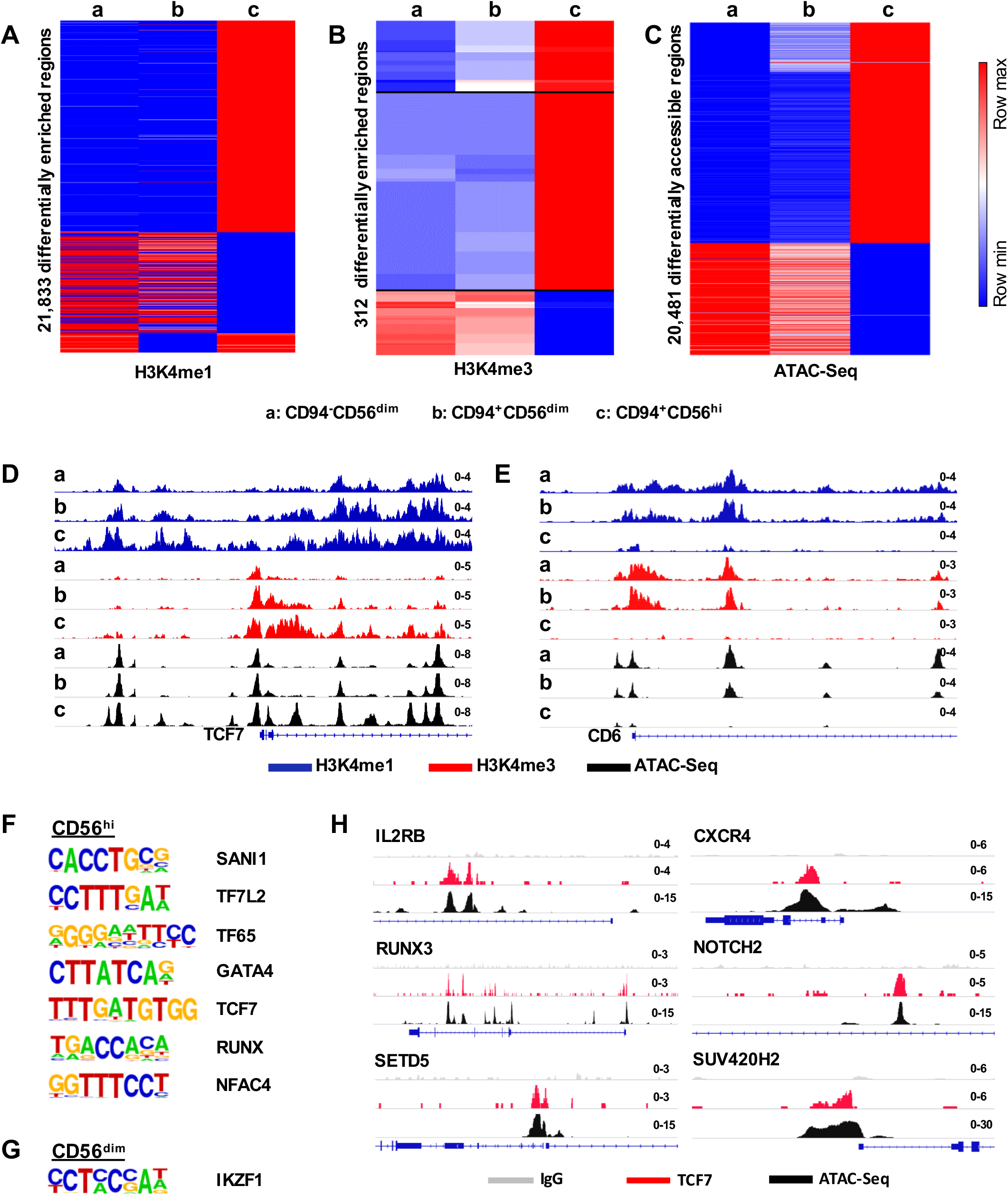
Distinct Chromatin Landscape in CD94^+^CD56^hi^ NK Cells. (A-C) Heatmap showing differential enrichment (>2 fold change of normalized counts) for H3K4me1 (A) and H3K4me3 (B) by CUT&RUN, or accessible chromatin by ATAC-Seq (C), in CD94^-^CD56^dim^, CD94^+^CD56^dim^, and CD94^+^CD56^hi^ NK cell subsets from 2 donors. (D, E) H3K4me1, H3K4me3, and ATAC-Seq signal on TCF7 (D) and CD6 (E), in the indicated NK cell subsets. (F, G) *De novo* analysis of transcription factor binding motifs enriched in open chromatin from CD56^hi^ NK cells (F), or CD56^dim^ NK cells (G), using HOMER. (H) ATAC-Seq and TCF7 CUT&RUN signal at the indicated loci from the three NK cell subsets. See also Figure S4.

### CD94^+^CD56^hi^ NK Cells Are *Bona Fide* Memory Lymphocytes

The CD94^+^CD56^hi^ NK cell transcriptome was enriched for transcripts associated with lymphocyte memory (Figure 3F; Table S5), including TCF7, CD44, SELL, CD70, IL7R, CCR1, CCR5, CCR7, BACH2, and DUSP4 (Baaten et al., 2010; Jeannet et al., 2010; Roychoudhuri et al., 2016; Weng et al., 2012). Chromatin accessibility and the density of H3K4me1 and H3K4me3 at these loci demonstrated that these genes were specifically remodeled for heightened expression in the CD94^+^CD56^hi^ NK subset (Figures 5D, 6A, and S5A). In contrast, the chromatin at genes typical of effector lymphocytes, including PRDM1, CD57, and KLRG1, (Brenchley et al., 2003; Kamimura and Lanier, 2015; Shin et al., 2013), was arranged for increased expression within CD56^dim^ NK cell subsets (Figures 6A and S5A). Consistent with the discrete developmental stages revealed by the analysis of chromatin, pairwise comparison of the three NK cell subsets showed that only CD94^+^CD56^hi^ NK cells were enriched for memory lymphocyte transcripts (Figures 6B and 6C).

**Figure 6.**
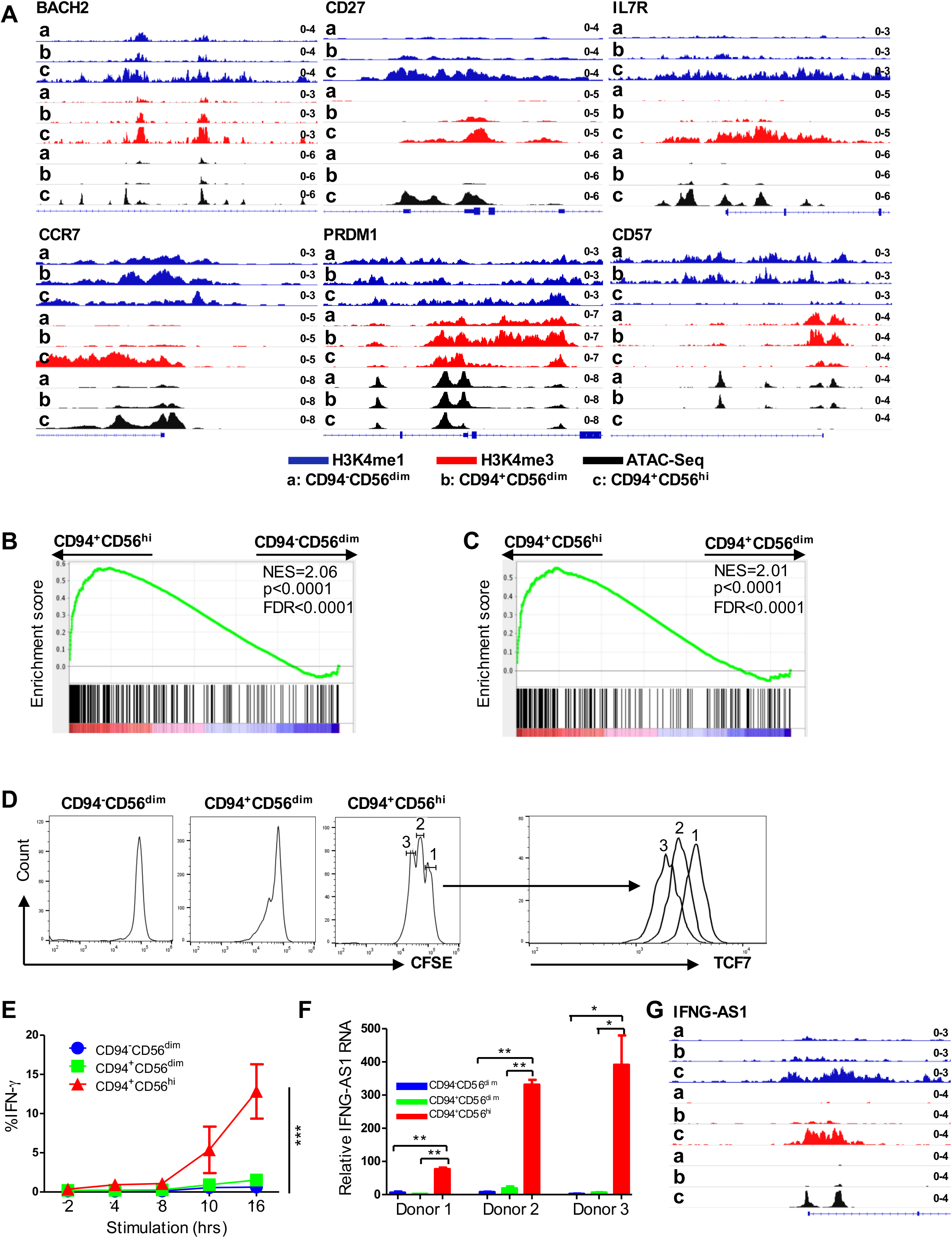
Chromatin, Transcriptome, and Phenotype Define CD94^+^CD56^hi^ NK Cells as Memory Cells. (A) Chromatin landscape by CUT&RUN and ATAC-Seq showing memory gene-associated loci in CD94^-^CD56^dim^, CD94^+^CD56^dim^, and CD94^+^CD56^hi^ NK cell subsets. Data is representative of 2 donors. (B, C) Gene set enrichment analysis comparing memory CD8^+^ T cell expression signature with CD94^+^CD56^hi^ and CD94^-^CD56^dim^ NK cells (B), or with CD94^+^CD56^hi^ and CD94^+^CD56^dim^ NK cells (C). (D) Sorted NK cell populations were labelled with CFSE to monitor proliferation and cultured for 5 days in IL-15 (5 ng/ml). CFSE and TCF7 were detected by flow cytometry. (E) Percent IFN-Y^+^ cells by flow cytometry after stimulation of sorted NK cell populations with IL-12 and IL-15 (n=4 donors); mean ± s.e.m.; two-way ANOVA, ***p<0.001. (F) IFNG-AS1 expression relative to GAPDH by RT-qPCR for the indicated NK cell populations sorted from 2 donors; two-tailed unpaired *t*-test, *p<0.05, **p<0.01. (G) IFNG-AS1 chromatin analysis as in (A). See also Figures S5 and S6.

As expected for memory T lymphocytes, stimulation of CD94^+^CD56^hi^ NK cells revealed a higher capacity to proliferate and produce IFN-γ than was observed with CD94^-^CD56^dim^ or CD94^+^CD56^dim^ NK cells (Figures 6D, 6E, and S5B). TCF7 was downregulated on sorted CD94^+^CD56^hi^ NK cells that had undergone multiple rounds of replication in response to stimulation (Figure 6D), as reported for the transition to effector cell phenotype that follows stimulation of memory CD8^+^ T cells (Lin et al., 2016a; Utzschneider et al., 2016). IFNG-AS1, a long noncoding RNA required for epigenetic activation of IFN-γ production (Gomez et al., 2013; Vigneau et al., 2003) was expressed from 30 to 200 times higher in CD94^+^CD56^hi^ NK cells than in the CD56^dim^ NK cell subsets (Figure 6F). The chromatin structure at the IFNG-AS1 locus (Figure 6G), as well as at other genes contributing to IFN-γ induction, including CD28, RUNX3, HLX, GZMB, and RAG2 (Figure S5) (Schoenborn and Wilson, 2007; Walker et al., 1999), indicate that robust IFN-γ production by CD94^-^CD56^dim^ NK cells in response to stimulation is a developmental adaptation characteristic of memory lymphocytes.

### WNT Signaling and TCF7 Are Required for Establishment of NK Cell Memory

To determine whether the WNT signaling and TCF7 activity associated with CD94^+^CD56^hi^ NK cells (Figures 3E-3H, 4G, 5D, 5F, 5H, S4D, and S6) are required for the establishment of memory, the effect of pharmacologic inhibition of WNT signaling and of TCF7 knockdown on their generation from sorted CD94^-^CD56^dim^ NK cells was assessed using established methods (Figure 7A and S7A) (Cooper et al., 2009; Romee et al., 2012). In response to stimulation with IL-12, IL-15, and IL-18 for 16 hrs, followed by 5 days in IL-15 alone, these cells upregulated CD94, TCF7, and IFNG-AS1 (Figure 7B-7D), and acquired a global transcriptional profile characteristic of memory T cells (Figures 7E and 7F). Upregulated genes typical of memory lymphocytes included DUSP4, CDKN1A, RGS1, CCR1, CCR5, IL2RA, CD70, CD74, ITGA1 and TNFSF10 (Figure 7F) (Henning et al. 2018; Weng et al. 2012; Hu and Chen 2013; O’Sullivan et al. 2015; Cerwenka and Lanier 2016). Downregulated genes characteristic of naive or effector lymphocytes included SELL, ITGA2, and KLRG1 (Figure 7F). All of these effects on stimulated CD94^-^CD56^dim^ NK cells were prevented if the WNT signaling inhibitor LGK974 was present in the culture (Figure 7B-7F). No toxicity was detected with LGK974 for up to eight days in culture, with or without stimulation, and the drug had no effect on CD94 levels in the absence of stimulation (Figure S7B). Specificity for the effect of LGK974 was demonstrated with M-110, an inhibitor of the NK cell kinase PIM3 that was highly enriched in TCF7^+^NK cells (Figure 3F); this drug had no effect on the cytokine-induced transition of CD94^-^NK cells to CD94^+^NK cells (Figure S7C). LGK974 had no effect on IFN-γ production after primary stimulation (Figure S7D), but the heightened IFN-γ production that was observed in response to secondary stimulation, after cells had transitioned to become CD94^+^CD56^high^ NK cells, was inhibited by LGK974 or by knockdown of TCF7 (Figures 7G, 7H, and S7E-S7G). These results demonstrate that TCF7 and WNT signaling are required for the establishment of CD94^+^CD56^high^ NK cells with a memory phenotype.

**Figure 7.**
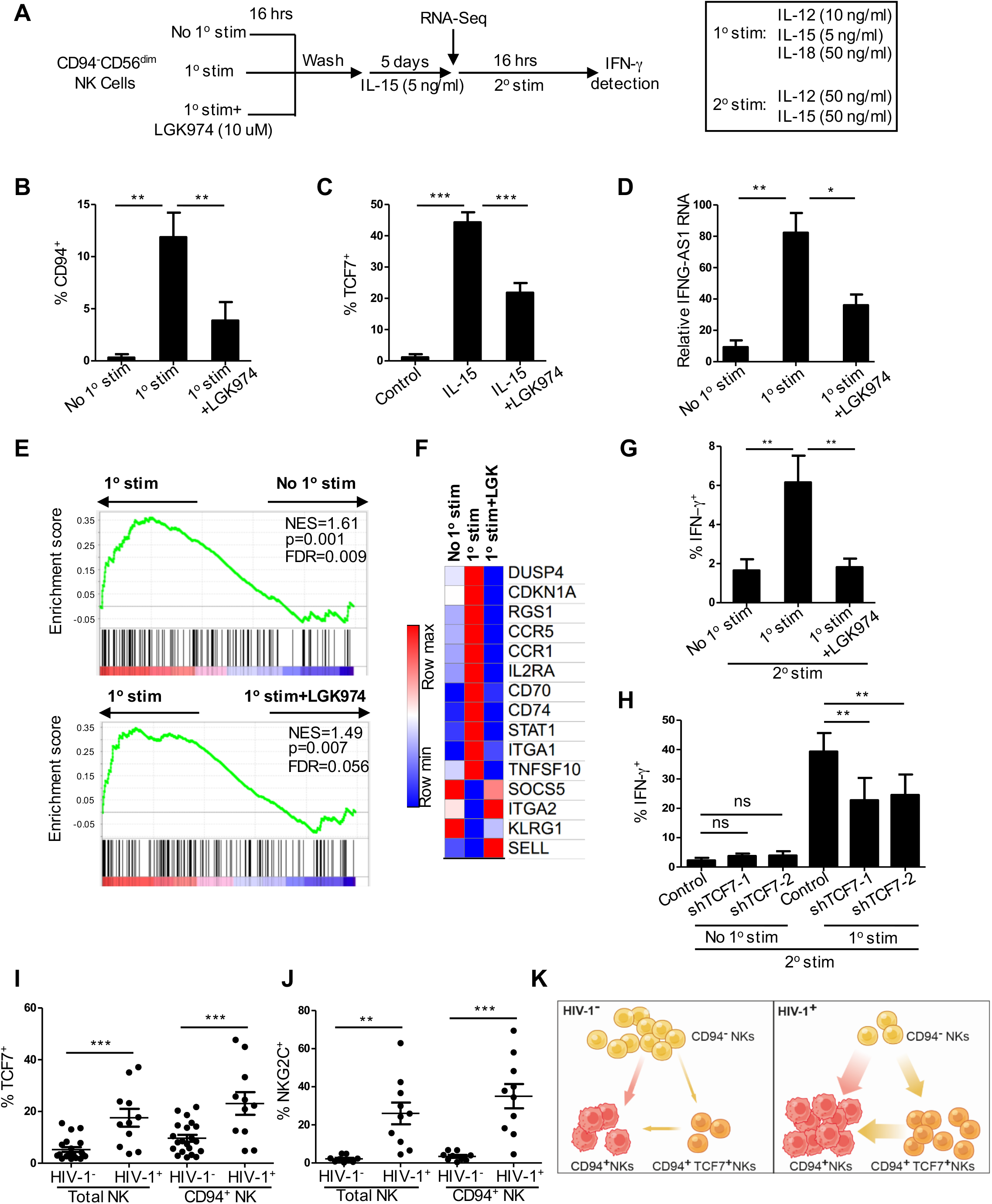
WNT Inhibition Blocks Cytokine Induced NK Memory Response. (A) Experimental scheme for *ex vivo* generation of memory NK cells using CD94^-^ CD56^dim^ NK cells sorted from blood. (B) CD94 by flow cytometry after 1° stimulation, with or without WNT inhibitor LGK974, as in (A), n=5. (C) TCF7 after culture in IL-15 for 5 days, with or without LGK974, n=5. (D) IFNG-AS1 expression relative to GAPDH by RT-qPCR after 1° stimulation and culture in IL-15 for 5 days, with or without LGK974, n=4. (E) Gene set enrichment analysis comparing memory CD8^+^ T cell expression signature with RNA-Seq data from sorted CD94^-^CD56^dim^ NK cells, with or without 1° stimulation and culture in IL-15 for 5 days, with or without LGK974. (F) Heatmap of memory related genes by RNA-Seq, in CD94^-^CD56^dim^ NK cells treated as indicated, with fold change of normalized counts Log2 >0.5, n=2. (G) Percent IFN-Y^+^ cells after 2° stimulation, following the indicated treatments, n=6. (H) Percent IFN-Y^+^ among GFP^+^ cells after transduction of Lin^-^CD56^+^ cells with lentivectors expressing GFP and shRNAs targeting either TCF7 or control, followed by stimulation, as indicated; n=3. (I) Percent TCF7^+^ among total NK cells (Lin^-^TBX21^+^) or CD94^+^NK cells (Lin^-^TBX21^+^CD94^+^) from HIV-1-(n=21) and HIV-1 ^+^ (n=11) individuals. (J) Percent NKG2C as in (I), from HIV-1^-^ (n=9) and HIV-1^+^ (n=10) individuals. (K) Model for effect of HIV-1 infection on NK cell subsets. Data is mean ± s.e.m.; two-tailed paired ř-test (B, C, D, G, and H); two-tailed unpaired ř-test (I and J); ns, not significant, *p<0.05, **p<0.01, ***p<0.001. See also Figure S7.

### Expansion of TCF7^+^ NK Cells in HIV-1 Infection

Finally, TCF7 was elevated when all Lin^-^TBX21^+^ cells in PBMCs from HIV-1^+^ people were examined (Figure 7I). The fact that TCF7 was also higher among the CD94^+^NK cell subset (Figure 7I) indicates that the TCF7 increase was independent of CD94 expression and suggests that the inflammation associated with HIV-1 infection drives the upregulation of TCF7^+^NK cells. A similar increase with HIV-1 infection was observed for another memory NK marker, NKG2C, consistent with reports that NKG2C^+^NK cells expand in response to SIV infection in rhesus macaques (Figures 7J and S7H) (Ram et al., 2018). Analogous to the expansion of memory T cells associated with the chronic inflammation that accompanies HIV-1 infection (Massanella et al., 2015), CD94^+^NK cells and CD94^+^TCF7^+^ memory NK cells are both likely to have expanded from CD94^-^ NK cells in response to the elevated systemic cytokines with HIV-1 infection (Figure 7K).

## DISCUSSION

Current antiretroviral drug regimens preserve CD4^+^ T cells and prevent progression to AIDS even when drug administration is delayed for years after HIV-1 infection (Günthard et al., 2016). Nonetheless, for reasons unknown, non-AIDS inflammatory pathology has been observed in such individuals (Deeks et al., 2013b). The experiments here demonstrate that ILCs and NK cells, either of which can be monitored in clinically-accessible tissues, are permanently disrupted by HIV-1 infection. These results offer explanation for HIV-1-associated inflammatory pathology, suggest new opportunities for prognosis of inflammatory disease in these patients, and open new windows on the physiology of these cells.

Inflammation associated with HIV-1 infection has been attributed to permanent disruption of the protective barrier posed by intestinal epithelium (Brenchley et al., 2006). The integrity of this tissue is maintained by homeostatic cytokines that are produced by lamina propria CD4^+^T_H_17 cells and ILC3s (Cella et al., 2009; Zheng et al., 2008). The experiments here demonstrate severe reduction in intestinal ILC3s sampled from HIV-1^+^ people on antiviral therapy with undetectable viremia and without significant reduction in lamina propria CD4^+^ T cells (Figure S1F). Given that blood ILCs are disrupted soon after HIV-1 infection (Kløverpris et al., 2016) it is likely that the abnormalities in the colon lamina propria ILCs are also established during acute HIV-1 infection, and that permanent loss of ILC3s, with attendant loss of epithelial barrier function, sustains chronic inflammation despite pharmacologic suppression of viremia. Given that ILC3s are accessible via lamina propria biopsy during screening colonoscopy (Figures 1C-1F), the likelihood of inflammatory complications might be predicted by assessing their numbers.

During acute infection with HIV-1, intestinal CD4^+^ T_H_17 cells are infected and killed. Though HIV-1 has been reported to infect ILCs (Zhao et al., 2018), these cells do not express CD4 or CCR5, the essential HIV-1 entry receptors, and infection of ILCs *ex vivo* cannot be detected with either CCR5-tropic or CXCR4-tropic HIV-1 (Figure S1G) (Kløverpris et al., 2016). These findings make it unlikely that ILCs are eliminated by direct HIV-1 infection. The effects of HIV-1 on ILC3s must therefore be indirect, resulting from JAK3 activation (Figures 1G and S1K) by the storm of common Y-chain cytokines that accompanies acute HIV-1 infection (Kacani et al., 1997; Roberts et al., 2010; Shebl et al., 2012; Stacey et al., 2009; Swaminathan et al., 2016).

In contrast to the CD4^+^ T cell recovery that is often observed with anti-HIV-1 therapy, disruption of ILCs appears to be permanent. Perhaps this is a consequence of the more primitive nature of ILCs, among which are included lymphoid tissue inducer cells necessary for establishment of secondary lymphoid tissue during embryonic development (Spits and Cupedo, 2012). If ILC depletion is an ongoing process, it might be possible to disrupt the cycle of inflammation with JAK3 inhibitors like CP-690550 used here to protect ILCs from cytokine stimulation *ex vivo* (Figure 1H).

CD94^+^NKG2C^+^NK cells are increased in HIV-1-infected people (Figures 2C and 2D and 7J) (Alter and Altfeld, 2009; Davis et al., 2016; Kottilil et al., 2004), but the total percentage of NK cells among PBMCs was not altered by HIV-1 (Figure 2A and 2B), demonstrating that CD94^+^NK cells increase at the expense of CD94^-^NK cells. Indeed CD94^-^NK cells stimulated *ex vivo* with inflammatory cytokines gave rise to CD94^+^NK cells (Figures 2C-2F and S2C-S2F), and pseudotime analysis of single cell transcriptomes from a range of NK cell types in the blood mapped a potential developmental trajectory from CD94^-^NK cells to CD94^+^NK cells. Stimulation of CD94^-^NK cells *ex vivo* gave rise to a relatively homogeneous population of CD94^+^TCF7^+^NK cells (Figure 3H). Taken together, the unbiased clustering of diverse NK cell transcriptomes, along with the *in vitro* differentiation assay, indicate that the increase in CD94^+^NK cells with HIV-1 infection result from a developmental shift in NK cells in response to stimulation by inflammatory cytokines (Figure 7K).

The studies presented here demonstrated that the subset of NK cells that is expanded with HIV-1 infection satisfies all standard criteria for *bona fide* memory cells. Pseudotime pathway analysis uncovered a gradient of TCF7 expression (Figures 3D and 3E), suggesting that this WNT transcription factor is responsible for driving the developmental transition of NK cells towards a memory state. TCF7^+^NK cells were found to overlap with CD56^hi^NK cells (Figure 3G), facilitating the subsequent isolation and molecular characterization of TCF7^+^NK cells (Figure 4A). As compared with other NK cell subsets, CD56^hi^NK cells had been reported to have increased proliferation rates and elevated cytokine production (Caligiuri, 2008; Poli et al., 2009), but the developmental implications of these phenotypes had not been clear. Global assessment of transcription, chromatin marks, and chromatin accessibility (Figures 4B-4E, 5A-5C, 6A, and S5A; Table S5), as well as proliferation capacity (Figure 6D), and the magnitude and kinetics of cytokine production in response to stimulation (Figures 6E), showed that, in all respects, these CD94^+^TCF7^+^CD56^hi^ NK cells resemble memory T cells, save the lack of clonotypic antigen receptors.

TCF7 is required for establishment and maintenance of memory T cells (Jeannet et al., 2010; Nish et al., 2017; Utzschneider et al., 2016). In a similar fashion, TCF7 was demonstrated here to be required for establishment and maintenance of memory NK cells. Analysis of the RNA-Seq datasets revealed TCF7/WNT as the dominant signaling pathway that distinguishes CD94^+^CD56^hi^ NK cells from other NK cell subsets (Figure 4G). CD94^-^CD56^dim^ cells upregulated TCF7, and other memory markers, after stimulation with IL-15 (Figure 3H). As determined by CUT&RUN, the TCF7 binding motif was enriched in CD94^+^CD56^hi^ NK cells, TCF7 bound preferentially to the open chromatin of genes associated with memory CD8^+^ T cells, and the chromatin of TCF7 target genes was more accessible by ATAC-Seq in CD94^+^CD56^hi^ NK cells (Figure 5F, 5H, S4D, and S6). A WNT inhibitor blocked the cytokine-induced transition of CD94^-^ CD56^dim^ NK cells to CD94^+^CD56^hi^ NK cells and the acquisition of memory cell-related features, including increased IFN-Y production and expression of molecules critical for survival and expansion, along with decreased expression of genes related to senescence and terminal differentiation (Figures 7B, 7C and 7E-7G). Finally, TCF7 knockdown inhibited memory-associated upregulation of IFN-Y production (Figure 7H). TCF7 may act as a transcription factor in the WNT signaling pathway or as an histone deacetylase (Xing et al., 2016). The fact that the TCF7 knockdown and the WNT-inhibitor had similar phenotypes (Figures 7G and H), as well as the results of the pathway analysis (Figure 4G), indicate that, by acting as a transcription factor in the WNT signaling pathway, TCF7 is an essential regulator of a real memory state in NK cells.

The expansion of TCF7^+^CD56^hi^NK cells in HIV-1^+^ individuals (Figure 7I-K) may occur secondarily to the inflammation that results from loss of gut-resident, homeostatic ILCs (Figure 1H). Consistent with this model, CD56^hi^NK cells are increased in systemic lupus erythematosus and Sjögren’s syndrome (Rusakiewicz et al., 2013; Schepis et al., 2009), two autoimmune diseases with chronic inflammation. If TCF7^+^CD56^hi^NK cells contribute to clinically significant pathology, the findings here may guide attempts to disrupt their numbers. For example, LGK974 (Figure 7C), a WNT inhibitor that is currently being tested in clinical trials (https://clinicaltrials.gov/ct2/show/NCT01351103), might provide clinical benefit by blocking the generation of TCF7^+^CD56^hi^NK cells from CD94^-^NK cells.

Conversely, the TCF7^+^NKG2C^+^CD94^+^CD56^hi^NK cells that are increased with HIV-1 infection (Figure 7J) may provide necessary antiviral protection, in which case, disruption of this NK cell subset would exacerbate the disease course in HIV-1 infection. People who inherit a deletion in the gene encoding NKG2C are at greater risk of HIV-1 infection, and have higher pretreatment viral loads and faster disease progression (Thomas et al., 2012). Specific KIR and HLA genotype combinations influence rates of HIV-1 acquisition and disease progression, perhaps due to KIR-mediated NK cell licencing or recognition of HIV-1-derived peptides by non-clonotypic receptors (Boulet et al., 2008; Lin et al., 2016b; Martin et al., 2002; Pelak et al., 2011). Interestingly, one patient with 30-50% CD56^hi^NK cells had nearly undetectable viral load and no opportunistic infections despite a CD4^+^ T cell count less than 50/mm^3^<50 (Fregni et al., 2013), suggesting that TCF7^+^NKG2C^+^CD94^+^CD56^hi^NK cells are critical to immune surveillance. NKG2C^+^NK cells are also increased with SIV infection of macaques (Ram et al., 2018) and this animal model may prove invaluable for clarifying the function of these memory NK cells. Better understanding of these memory NK cells may lead to new approaches for preventing and controlling HIV-1 infection, other infections, or autoimmune disease.

## METHODS

### Clinical samples

All human blood and colon samples were collected from participants who had provided written informed consent for protocols that included study of cellular immunity in HIV-1 infection, in accordance with procedures approved by the University of Massachusetts Medical School (UMMS) Institutional Review Board. Routine screening colonoscopy was scheduled as medically indicated. HIV-1^-^ control individuals undergoing colonoscopy the same day were matched for gender and age. In pilot experiments with two patients, multiple biopsies of different segments of the bowel from the same patient were compared and no differences in ILC populations were found. Subsequent sampling was from the descending colon only. All patients were consented the day of the procedure. All blood and colon samples that were collected and processed were included in the data analysis after simple allocation into one of two groups, HIV-1^-^ or HIV-1^+^. Attempts to stratify samples according to viral load or CD4^+^ T cell count of the donor failed to detect significant differences between the subgroups (Figures S1 and S2).

### Human mononuclear cell isolation

Human peripheral blood or umbilical cord blood was diluted with an equal volume of RPMI-1640 (Gibco), overlaid on Histopaque (Sigma), and centrifuged at 500 x g for 30 mins at room temperature. Mononuclear cells were washed 3 times with MACS buffer (0.5% BSA and 2 mM EDTA in PBS) and either used immediately or frozen in FBS containing 10% DMSO.

### Human intestinal lamina propria lymphoid cell (LPL) preparation

Human intestinal biopsies were incubated with PBS containing 5 mM EDTA, 150 μg/ml DTT, and 0.1% β-mercaptoethanol on a shaker for 15 min at 37^-^ to remove epithelial cells. The remaining tissue containing the LPLs was washed with complete RPMI-1640 (10% FBS, 1:100 GlutaMAX, and 1:1000 β-mercaptoethanol), then digested with RPMI-1640 containing 125 μg/ml Liberase (Roche) and 200 μg/ml DNase I (Sigma) for 30 min on a shaker at 37^-^C LPLs were filtered through a 70 μm cell strainer and washed with RPMI-1640.

### Definition of human lineage-negative lymphoid populations

Live singlets of human PBMCs, or of intestinal lamina propria lymphoid cells, were stained with a panel of lineage markers (Lin): anti-CD3, anti-TCRαβ, anti-CD19, anti-B220, anti-CD34, anti-FcsRIa, anti-CD11c, anti-CD303, anti-CD123, anti-CD1a, anti-CD14, and anti-CD94. The specific antibodies used are listed in Table S5. Cells that were negative for all lineage markers were defined as lineage-negative (Lin^-^) lymphoid cells.

### Human NK cell enrichment

NK cells were negatively enriched by excluding T cells, B cells, stem cells, dendritic cells, monocytes, granulocytes, and erythroid cells with the human NK isolation kit (MACS) according to manufacturer's instruction.

### Flow cytometry

Live and dead cells were discriminated using the Live and Dead violet viability kit (Invitrogen, L-34963). For cell surface molecule detection, the cells were resuspended in antibody-containing MACS buffer for 30-60 min at 4°C in the dark. To detect cytokine production, cells were stimulated with the indicated cytokines for 16 hrs, or with PMA and ionomycin (cell stimulation cocktail 00-4970-03, ebioscience) for 3-6 hrs. In both cases, protein transport inhibitor (00-4980-03, eBioscience) was present during the stimulation. For intracellular staining of transcription factors or cytokines, cells were fixed and permeabilized using Foxp3 staining buffer kit (eBioscience) and target intracellular molecules were stained as for surface staining.

### Cell sorting

PBMCs were stained with a panel of lineage markers that included anti-CD3, anti-TCRαβ, anti-CD19, anti-B220, anti-CD34, anti-FcsRIa, anti-CD11c, anti-CD303, anti-CD123, anti-CD1a, and anti-CD14, The Lin^-^population was sorted based on cell surface CD56 and CD94, as indicated in the legend to Figures S2G and S4A using a BD FACSAria IIu

### Degranulation assay

PBMCs were seeded in a 24 well plate at 2 × 10^6^ cells/well in RPMI-1640 with anti-CD107a antibody (Biolegend, 1:200). Then, cells were treated with PMA/ionomycin and protein transport inhibitor for 5 hrs in a 37°C incubator with 5% CO_2_. Surface CD107a was detected by flow cytometry.

### Killing assay

K562 or Jurkat cells were washed and resuspended at 10^6^ cell/ml in PBS. Calcein-AM (Neri et al., 2001) (Molecular Probes) was added at 10 μM and the cells were labelled at 37°C in 5% CO_2_ for 30 min. After washing with PBS for 2 times, the labelled cells were resuspended in complete RPMI-1640 and aliquoted in a V bottom 96 well plate (5,000 cells in 100 μl). The sorted NK cells were resuspended in complete RPMI-1640 and the concentration was adjusted at 15 fold of target cells (7.5×10^4^ cells) in 50μl. The effector and target cells were mixed and centrifuged at 50 x g for 0.5 min and were incubated at 37°C in 5% CO_2_ for 6 hrs. Cells were pelleted and 100 μl supernatant was transferred to a 96-well solid black microplate. The fluorescence released by labelled target cells were detected using BioTek Synergy 2 plate reader (excitation filter: 485/20 nm; band-pass filter: 528/20 nm). Specific lysis was determined as: [(test fluorescence release – spontaneous fluorescence release) / (maximum fluorescence release – spontaneous fluorescence release)] × 100.

### Lentivirus production and TCF7 knockdown

TCF7-specific pGIPZ shRNAs (Dharmacon), were subcloned into a modified pAGM plasmid and this was used to transfect HEK293 cells to generate VSV G pseudotyped lentivirus, as described (Pertel et al., 2011). Lin^-^CD56^+^ NK cells were subjected to 3 rounds of transduction before 1° stimulation.

### Cytokine induced NK cell memory assay

PBMCs or sorted CD94^-^NK cells were prestimulated with IL-15 (5 ng/ml) ^+^ IL-12 (10 ng/ml) ^+^ IL-18 (50 ng/ml) for 16 hrs, or cells were incubated with IL-15 alone (5 ng/ml) to maintain cell survival (no pre-stimulation, control). Cytokines were washed out and the cells were rested by incubation in IL-15 (5 ng/ml) alone for 5 days. Cells were then re-stimulated with IL-12 (50 ng/ml) ^+^ IL-15 (50 ng/ml) for 16 hrs. IFN-γ production was detected by intracellular staining with anti-IFN-Y antibody using a Foxp3 staining buffer kit (eBioscience). As compared with cells that had not been pre-stimulated, the increase in IFN-γ production in pre-stimulated cells was used as a readout for pre-stimulation induced memory, as previously described (Poli et al., 2009).

### Library preparation for bulk RNA-Seq

The bulk RNA sequencing library was prepared using CEL-Seq2 (Hashimshony et al., 2016). RNA of sorted cells was extracted using TRIzol reagent. 10 ng RNA was used for first strand cDNA synthesis using barcoded primers containing unique molecular identifier (UMI) sequences. Specifically, for CD94^-^NK cell and CD94^+^NK cell 2 population RNA-Seq from 2 donors, the RNA of CD94^-^NK cells was reverse transcribed with 5’-GCC GGT AAT ACG ACT CAC TAT AGG GAG TTC TAC AGT CCG ACG ATC NNN NNN AGA CTC TTT TTT TTT TTT TTT TTT TTT TTT V-3’ and 5’-GCC GGT AAT ACG ACT CAC TAT AGG GAG TTC TAC AGT CCG ACG ATC NNN NNN CAG ATC TTT TTT TTT TTT TTT TTT TTT TTT V-3’. The RNA of CD94^+^NK cells was reverse transcribed with 5’-GCC GGT AAT ACG ACT CAC TAT AGG GAG TTC TAC AGT CCG ACG ATC NNN NNN CAT GAG TTT TTT TTT TTT TTT TTT TTT TTT V-3’ and 5’-GCC GGT AAT ACG ACT CAC TAT AGG GAG TTC TAC AGT CCG ACG ATC NNN NNN TCA CAG TTT TTT TTT TTT TTT TTT TTT TTT V-3’. The second strand was synthesized using NEBNext Second Strand Synthesis Module (NEB). The pooled dsDNA was purified with AMPure XP beads (Beckman Coulter, A63880) and subjected to in vitro transcription (IVT) using HiScribe T7 High Yield RNA Synthesis Kit (NEB), then treated with ExoSAP-IT (Affymetrix 78200). IVT RNA was fragmented using RNA fragmentation reagents (Ambion) and underwent a 2^nd^ reverse transcription step using random hexamer RT primer-5’-GCC TTG GCA CCC GAG AAT TCC ANN NNN N-3’ to incorporate the second adapter. The final library was amplified with indexed primers: RP1-5’-AAT GAT ACG GCG ACC ACC GAG ATC TAC ACG TTC AGA GTT CTA CAG TCC GA-3’ and RPI1-5’-CAA GCA GAA GAC GGC ATA CGA GAT CGT GAT GTG ACT GGA GTT CCT TGG CAC CCG AGA ATT CCA-3’ and the purified library was quantified with 4200 TapeStation (Agilent Technologies) and paired-end sequenced on a Nextseq 500 V2 (Illumina) using 15 cycles Read 1, 6 cycles Index 1, and 71 cycles Read 2.

RNA-Seq on CD94^-^CD56^dim^, CD94^+^CD56^dim^ and CD94^+^CD56^hi^ NK cells from 4 donors was performed as above. Specifically, the RNA of CD94^-^CD56^dim^ NK cells was reverse transcribed with 5’-GCC GGT AAT ACG ACT CAC TAT AGG GAG TTC TAC AGT CCG ACG ATC NNN NNN AGA CTC TTT TTT TTT TTT TTT TTT TTT TTT V-3’; 5’-GCC GGT AAT ACG ACT CAC TAT AGG GAG TTC TAC AGT CCG ACG ATC NNN NNN TCA CAG TTT TTT TTT TTT TTT TTT TTT TTT V-3’; 5’-GCC GGT AAT ACG ACT CAC TAT AGG GAG TTC TAC AGT CCG ACG ATC NNN NNN ACC ATG TTT TTT TTT TTT TTT TTT TTT TTT V-3’; and 5’-GCC GGT AAT ACG ACT CAC TAT AGG GAG TTC TAC AGT CCG ACG ATC NNN NNN CTG TGA TTT TTT TTT TTT TTT TTT TTT TTT V-3’. CD94^+^CD56^dim^ cells were reverse transcribed with 5’-GCC GGT AAT ACG ACT CAC TAT AGG GAG TTC TAC AGT CCG ACG ATC NNN NNN CAT GAG TTT TTT TTT TTT TTT TTT TTT TTT V-3’; 5’-GCC GGT AAT ACG ACT CAC TAT AGG GAG TTC TAC AGT CCG ACG ATC NNN NNN GTC TAG TTT TTT TTT TTT TTT TTT TTT TTT V-3’; 5’-GCC GGT AAT ACG ACT CAC TAT AGG GAG TTC TAC AGT CCG ACG ATC NNN NNN ACT CGA TTT TTT TTT TTT TTT TTT TTT TTT V-3’ and 5’-GCC GGT AAT ACG ACT CAC TAT AGG GAG TTC TAC AGT CCG ACG ATC NNN NNN TGC AGA TTT TTT TTT TTT TTT TTT TTT TTT V-3’. CD94^+^CD56^hi^ NK cells were reverse transcribed with 5’-GCC GGT AAT ACG ACT CAC TAT AGG GAG TTC TAC AGT CCG ACG ATC NNN NNN CAG ATC TTT TTT TTT TTT TTT TTT TTT TTT V-3’; 5’-GCC GGT AAT ACG ACT CAC TAT AGG GAG TTC TAC AGT CCG ACG ATC NNN NNN GTT GCA TTT TTT TTT TTT TTT TTT TTT TTT V-3’; 5’-GCC GGT AAT ACG ACT CAC TAT AGG GAG TTC TAC AGT CCG ACG ATC NNN NNN ACG TAC TTT TTT TTT TTT TTT TTT TTT TTT V-3’ and 5’-GCC GGT AAT ACG ACT CAC TAT AGG GAG TTC TAC AGT CCG ACG ATC NNN NNN CAA CCA TTT TTT TTT TTT TTT TTT TTT TTT V-3’.

For sequencing of CD94^-^CD56^dim^ NK cells in the cytokine-induced NK cell memory model, RNA collected after no 1° stimulation, after 1° stimulation, and after 1° stimulation in the presence of LGK974, was reverse transcribed with 5’-GCC GGT AAT ACG ACT CAC TAT AGG GAG TTC TAC AGT CCG ACG ATC NNN NNN GTC TAG TTT TTT TTT TTT TTT TTT TTT TTT V-3’, 5’-GCC GGT AAT ACG ACT CAC TAT AGG GAG TTC TAC AGT CCG ACG ATC NNN NNN GTT GCA TTT TTT TTT TTT TTT TTT TTT TTT V-3’ and 5’-GCC GGT AAT ACG ACT CAC TAT AGG GAG TTC TAC AGT CCG ACG ATC NNN NNN ACC ATG TTT TTT TTT TTT TTT TTT TTT TTT V-3’ respectively, The final library was amplified with indexed primers: RP1-5’-AAT GAT ACG GCG ACC ACC GAG ATC TAC ACG TTC AGA GTT CTA CAG TCC GA-3’ and RPI1 (for donor 1) −5’-CAA GCA GAA GAC GGC ATA CGA GAT CGT GAT GTG ACT GGA GTT CCT TGG CAC CCG AGA ATT CCA-3’ or RPI2 (for donor 2)-5’-CAA GCA GAA GAC GGC ATA CGA GAT ACA TCG GTG ACT GGA GTT CCT TGG CAC CCG AGA ATT CCA-3’.

### Library preparation for single cell RNA-Seq

The single cell sequencing library was prepared using InDrop barcoding (Klein et al., 2015). Individual, pre-sorted cells were captured in droplets containing lysis buffer, reverse transcription reagents, and hydrogel microspheres carrying primers for barcoding, each captured cell using a custom-built InDrop microfluidics system, as described (Klein et al., 2015). After cDNA for each cell was synthesized in individual droplets with unique barcodes, the droplet emulsion was broken and cDNA from all cells was pooled. A single cell cDNA library was prepared using essentially CEL-Seq2 (Hashimshony et al., 2016) as described above for bulk RNA-Seq, with the following changes. PE2-N6-5’-TCG GCA TTC CTG CTG AAC CGC TCT TCC GAT CTN NNN NN-3’ was used in the 2^nd^ reverse transcription reaction. CD94^-^NK cells libraries were linearly amplified with Indexed primers PE1-6: 5’-CAA GCA GAA GAC GGC ATA CGA GAT ATT GGC CTC TTT CCC TAC ACG A-3’ or PE1-17: 5’-CAA GCA GAA GAC GGC ATA CGA GAT CTC TAC CTC TTT CCC TAC ACG A-3’ and PE2: 5’-AAT GAT ACG GCG ACC ACC GAG ATC TAC ACG GTC TCG GCA TTC CTG CTG AAC-3’. Libraries from CD94^+^NK cells were amplified with index 1 primers PE1-12: 5’-CAA GCA GAA GAC GGC ATA CGA GAT TAC AAG CTC TTT CCC TAC ACG A-3’ or PE1-18: 5’-CAA GCA GAA GAC GGC ATA CGA GAT GCG GAC CTC TTT CCC TAC ACG A-3’, and PE2 (above). Libraries were sequenced 38 cycles Read 1, 6 cycles Index 1, 48 cycles Read 2 using Illumina Nextseq 500 V2 with custom sequencing primers added to the cartridge: R1 5’-GGC ATT CCT GCT GAA CCG CTC TTC CGA TCT-3’ Idx: 5’-AGA TCG GAA GAG CGT CGT GTA GGG AAA GAG-3’ R2 5’-CTC TTT CCC TAC ACG ACG CTC TTC CGA TCT-3’.

### CUT&RUN

Sorted CD94^-^CD56^dim^, CD94^+^CD56^dim^ and CD94^+^CD56^hi^ NK cells were processed as described in previously published papers (Hainer et al., 2018; Skene and Henikoff, 2017; Skene et al., 2018). Briefly, cells were lysed in nuclear extraction buffer, the crude nuclei were precipitated by centrifuge then were resuspended and bind with Bio-Mag Plus Concanavalin A coated beads (Polysciences, cat# 86057), after blocking for 5 min, primary antibodies were added and incubated at 4 °C overnight, then pA-MNase was added and incubate for 1 hr at 4 °C, CaCl_2_ was added to activate digestion, after 30 min, stop buffer was added, the supernatant containing released chromatin was subject to pheno-chloroform-isoamyl extraction. The sequencing library was constructed according to NEBNext Ultra II DNA library Prep kit for Illumina (NEB, cat#7645L), NEBNext Multiplex Oligos for Illumina (NEB, cat# E6609S). For donor 1, sorted CD94^-^ CD56^dim^ NK cells were treated with Rabbit anti-human IgG (Control, abcam, cat# ab2410), Rabbit anti-H3K4me1 (diagnode, cat# c15410194), Rabbit anti-H3K27ac (diagnode, cat# c15410196), Rabbit anti-H3K4me3 (diagnode, cat# c15410003), Rabbit anti-TCF7 (cell signaling technology, cat# 2203S), Rabbit anti-H3K27me3 (diagnode, cat# c15410195), primers D1-D6 were used for library amplification for each antibody treated sample according to above antibody order. Accordingly, primers D7-D12 were used for sorted CD94^+^CD56^dim^ NK cells related samples and primers E1-E6 were used for sorted CD94^+^CD56^hi^ NK cells related samples. For donor 2, primers E7-E12 were used for sorted CD94^-^CD56^dim^ NK cells, primers F1-F6 were used for sorted CD94^+^CD56^dim^ NK cells, and primers F7-F12 were used for sorted CD94^+^CD56^hi^ NK cells. Paired-end sequenced on a Nextseq 500 V2 (Illumina) using 45 cycles Read 1, 8 cycles Index 1, and 32 cycles Read 2.

### ATAC-Seq

Sorted CD94^-^CD56^dim^, CD94^+^CD56^dim^ and CD94^+^CD56^hi^ NK cells were processed as described in previous published papers (Buenrostro et al., 2013, 2015). Briefly, crude nuclei was precipitated after cell lysis, and treated with Tn5 transposase (Nextera DNA Library Prep Kit (Illumina, cat# FC-121-1030) for 30 min, released DNA fragments were purified, then were subjected to library preparation. Barcode primers used for CD94^-^CD56^dim^, CD94^+^CD56^dim^ and CD94^+^CD56^hi^ NK cells from donor 1 library construction were: 5’-CAA GCA GAA GAC GGC ATA CGA GAT TCG CCT TAG TCT CGT GGG CTC GGA GAT GT-3’, 5’-CAA GCA GAA GAC GGC ATA CGA GAT CTA GTA CGG TCT CGT GGG CTC GGA GAT GT-3’ and 5’-CAA GCA GAA GAC GGC ATA CGA GAT TTC TGC CTG TCT CGT GGG CTC GGA GAT GT-3’ respectively. Barcode primers used for CD94^-^CD56^dim^, CD94^+^CD56^dim^ and CD94^+^CD56^hi^ NK cells from donor 2 library construction were: 5’-CAA GCA GAA GAC GGC ATA CGA GAT GCT CAG GAG TCT CGT GGG CTC GGA GAT GT-3’, 5’-CAA GCA GAA GAC GGC ATA CGA GAT AGG AGT CCG TCT CGT GGG CTC GGA GAT GT-3’ and 5’-CAA GCA GAA GAC GGC ATA CGA GAT CAT GCC TAG TCT CGT GGG CTC GGA GAT GT-3’ respectively. The common primer sequence was: 5’-AAT GAT CGG CGA CCA CCG ATA TCT ACA CTC GTC GGC AGC GTC AGA TGTG-3’. Paired-end sequenced on a Nextseq 500 V2 (Illumina) using 42 cycles Read 1, 8 cycles Index 1, and 42 cycles Read 2.

### Bulk RNA-Seq Processing and Analysis

The pooled sets of RNA-Seq reads were separated by CEL-Seq barcodes and mapped to the HG19 genome using Tophat (Kim et al., 2013) (version 2.0.14, default parameters). Aligned reads were quantified by ESAT (Derr et al., 2016) using a transcript annotation file containing all RefSeq genes filtered to select only ‘NM’ transcripts and extending the transcripts up to 1,000 bases past the end of the annotated 3’ end *(-wExt 1000, -task score3p)*, discarding multimapped reads *(-multimap ignore)*. The most varied genes were analyzed using DEBrowser, differential expression analysis was performed using DESeq2 (Love et al., 2014), and MA plots were created using values from lfcShrink within DESeq2. Any gene differentially expressed in these 3 tests in Figure 4B was shown. For principal component analysis (PCA), data was transformed using rlog within DEseq2 and prcomp was used to calculate the PCs.

### Single cell RNA-Seq processing and analysis

Reads were mapped to the HG19 genome using Tophat (Kim et al., 2013) (version 2.0.14, default parameters). To assign each read to a given cell and collapse duplicate reads into single UMIs, alignments were processed by ESAT (Derr et al., 2016) using its single-cell analysis module (-*scPrep)* with the same transcript annotation file used for the bulk RNA-Seq analysis, extending the transcripts up to 1,000 bases past the annotated 3’ end *(-wExt 1000, – task score3p)*, discarding multimapped reads *(-multimap ignore)*, and requiring a single observation of a UMI for a transcript to be counted *(-umiMin* 1).

The final output of ESAT is a table where rows are genes, cells are columns, and values represent the number of UMIs detected in each cell. The dataset was loaded into R and, if not part of base R, the packages used are noted. First, data was normalized using the TMM method from EdgeR (Robinson and Oshlack, 2010). On this matrix, PCA was run in order to determine the number of dimensions that contribute variance to the data and to select genes that were highly variable in the dataset. ICA was run using the fastICA (Hyvärinen and Oja, 2000) algorithm to reduce the data to the number of dimensions that contained variance. To determine the optimum number of cell clusters in the dataset the methods of (Tibshirani and Walther, 2005) were utilized. This analysis showed 2 clusters gives the highest predictive strength, and that further subdividing of the cells into additional clusters severely lowered the score. Spectral clustering was then run with n of 2 centers and the symmetrical method on the cell’s ICA components using the kknn package (Hechenbichler and Schliep, 2004). Differential expression analysis was used to determine the genes that were most significantly different between the two clusters (Robinson et al., 2010). For visualization, the cells were then reduced to 2 dimensions using the Rtsne package, which took the ICA components as input (Van Der Maaten, 2014). Lastly, the minimum spanning tree (MST) was constructed on the tSNE plot utilizing Monocle (Trapnell et al., 2014). For pseudotime analysis, Monocle’s built-in differential expression tools were utilized. For visualization purposes, tSNE mapped cells were color coded by the expression of given genes; a weighted density map was created that takes into account both the number of cells in a region as well as their expression values. Heatmaps were generated using the list of genes that were found to be significantly differentially expressed in both the spectral clustering clusters and with the pseudotime analysis. Rows of the heatmap were grouped by similarity (heatmap.2), and columns were ordered based on the ordering provided by the pseudotime analysis.

### CUT&RUN and ATAC-Seq data analysis

Data processing made using in-house tool called dolphin (https://dolphin.umassmed.edu/). For CUT&RUN, paired-end reads were removed where the average quality scores in window size 10 are less then 15 and trimmed where leading and trailing bases with quality scores less than 15 using trimmomatic version 0.32. Reads that were longer than 25 bases after trimming were kept for further analysis. The reads were then aligned to human reference genome hg19 using Bowtie2 with options -un-conc to filter out reads that align un-concordantly. Duplicated reads were filtered out using Picard’s MarkDuplicates version 0.32. Peaks were then called using MACS2 (Zhang et al., 2008). Alignment files were also converted to tdf format using IGVtools count function version 2.3.31 using -w 5 parameter. For ATAC-Seq, first, paired-end reads were filtered which have bed qualities using trimmomatic version 0.32, and then aligned with Bowtie2, version 2.2.3 to a reference genome hg19. The duplicate alignments were then removed using Picard’s MarkDuplicates version 0.32. To be able to accurately call the peaks. each aligned read was first trimmed to the 9-bases at the 5’ end, the region where the Tn5 transposase cuts the DNA in the performed ATAC-Seq experiment. To smooth the peaks the start site of the trimmed read extended 10-bases up and down stream. Peaks were called using these adjusted aligned reads with MACS2 (Zhang et al., 2008). For visualization, the adjusted aligned reads were converted to tdf files using IGVTools, version 2.3.31 (Robinson et al., 2011) (IGVtools count -w 5). For motif analysis, motifs enriched in accessible regions within the enhancers and promoters of indicated NK subset was found by using findMotifsGenome from HOMER2 package using the accessible regions within the enhancers and promoters of the other two NK subsets as background. The de-novo motifs found were matched against HOCOMOCCO11 motif data base.

### Gene set enrichment analysis

The RNK file including gene name and log2 fold change, comparing CD94^+^CD56^hi^ vs CD94^-^CD56^dim^ and CD94^+^CD56^hi^ vs CD94^+^CD56^dim^, were used as input for GenePattern (https://genepattern.broadinstitute.org/gp/pages/index.isf). The gene set database selected was C7 (immunologic signatures) and the number of permutations was set to 5000.

## QUANTIFICATION AND STATISTICAL ANALYSIS

Statistical analysis was performed with GraphPad Prism software using paired or unpaired two-tailed student’s *t*-test or Mann Whitney test as indicated in the figure legends. p<0.05 was considered as significant. Variance was estimated by calculating the mean ± s.e.m. in each group. Variances among groups of samples were compared using the F-test function in GraphPad.

## DATA AND SOFTWARE AVAILABILITY

### Bulk and single-cell RNA-Seq, CUT&RUN, and ATAC-Seq datasets

Datasets can be found under SuperSeries GSE122326 at https://www.ncbi.nlm.nih.gov/geo/query/acc.cgi?acc=GSE97727.

GSE97727: CD94^-^ and CD94^+^ NK cell bulk and single cell RNA-Seq GSE122324: CD94^-^CD56^dim^, CD94^+^CD56^dim^, and CD94^+^CD56^hi^ NK cells RNA-Seq GSE122325: CD94^-^CD56^dim^ NK cells, 1° stim and 5 day culture RNA-Seq GSE122548: CD94^-^CD56^dim^, CD94^+^CD56^dim^, and CD94^+^CD56^hi^ NK cells ATAC-Seq GSE122549: CD94^-^CD56^dim^, CD94^+^CD56^dim^, and CD94^+^CD56^hi^ NK cells CUT&RUN

## SUPPLEMENTAL INFORMATION

Supplemental information includes 7 figures and 5 tables.

## ACKNOWLEDGEMENTS

We thank the patients who generously provided blood and colon biopsy samples, as well as their caretakers, Drs. Jennifer Daly, Sarah Cheeseman, and Mireya Wessolossky of the University of Massachusetts Medical School (UMMS). Caitlin Mannarino, Anne Foley, and Margaret McManus (UMMS) provided IRB regulatory assistance, sample preparation, and record keeping. Katherine Luzuriaga (UMMS) supported the patient sample database and repository. Alex Ratner, Sarah Boswell, and Allon Klein (Harvard Medical School) contributed technical assistance and barcoded hydrogel beads. Thomas Fazzio and Tong Wu provided technical support and protein A-MNase for CUT&RUN. David Artis, Leslie Berg, Marco Colonna, Jun Huh, Joonsoo Kang, and Susan Swain offered invaluable advice. This research was supported by NIH U01HG007910 (to M.G. and J.L.), R01AI111809 (to J.L.), DP1DA034990 (to J.L.), R21AI119885 (to M.G.), R01DK105837 (to M.G.), and the Translational Medicine Core of the University of Massachusetts Center for AIDS Research (P30AI042845). Requests for resources and reagents should be directed to Jeremy Luban.

## AUTHOR CONTRIBUTIONS

Y.W. and J.L. designed the experiments. Y.W. performed the experiments with assistance from S.J., K.G., A.D., L.L., S.M., K.K., P.V. and P.M. Y.W. and J.L. analyzed the experimental data. Y.W., K.G., A.D. P.V., L.L., A.K., M.G., and J.L. analyzed the expression data. T.G. and J-M.H. obtained and provided clinical samples. Y.W. and J.L. wrote the manuscript which was revised and approved by all authors.

