## Supplemental Figures and Tables for "HIV-1-induced cytokines deplete homeostatic ILCs and expand TCF7-dependent memory NK cells"

7 Supplementary Figures with Legends

5 Supplementary Tables,

### SUPPLEMENTARY FIGURE LEGENDS

**Figure S1. ILC Gating and Effect of HIV-1. Related to Figure 1.**

(A) Flow cytometry gating on lymphoid, singlet, live PBMCs, and on lineage markers as indicated on the x-axis. The y-axis shows empty APC channel without antibody staining.

(B) PBMCs gated on 12 lineage markers (y axis) as in (A) versus CD127 on the x axis. $Lin^-$ cells were stained for TBX21, CRTH2, or RORγT, or stained for IFN-γ, IL-13, or IL-22 after 3 hrs stimulation with PMA/ionomycin.

(C) Percent $CD127^+$ILCs among $Lin^-$PBMCs from people who are HIV-$1^-$, n=23; HIV-$1^+$ with viremia $>10^4$/ml (HIV-$1^{high}$), n=11; or HIV-$1^+$ with viremia $<10^2$/ml (HIV-$1^{low}$), n=11.

(D) Percent $CD127^+$ILCs among $Lin^-$PBMCs from individuals who are HIV-$1^-$, n=23; HIV-$1^+$ with $CD4^+$T cells $>500/mm^3$, n=7; or HIV-$1^+$ with $CD4^+$T cells $<500/mm^3$, n=10.

(E) Colon lamina propria lymphoid cells were stimulated as in (B) and $Lin^-$ cells were stained for IL-22 and CD127.

(F) Percent $CD4^+$T cells among colon lamina propria lymphocytes from HIV-$1^-$ (n=3) and HIV-$1^+$ (n=5) individuals.

(G) The trimmed mean of M-values normalization method (from DEBrowser) for the indicated genes, based on RNA-Seq data from sorted $Lin^-CD127^+$ILCs (n=4 PBMC donors).

(H-K) PBMCs were treated with the indicated cytokines (H and K, n=4), chemokines (I, n≥2), TLR agonists (J, n=4), or L-kynurenine (H) for 16 hrs. Percent $CD127^+$ among $Lin^-$ cells is shown.

Data is mean ± s.e.m.; two tailed unpaired t-test; ns, non-significant, $^{*}p<0.05$, $^{**}p<0.01$, $^{***}p<0.001$.

**Figure S2. HIV-1 Infection Increases $CD94^+$NK cells. Related to Figure 2.**

(A, B) $Lin^-TBX21^+$ PBMCs were stained for CD56 (A), and for EOMES and CD94 (B).

(C) Percent $Lin^-TBX21^+$ PBMCs from HIV-$1^-$ (n=14) and HIV-$1^+$ (n=22) individuals (lineage markers include anti-CD94).

(D) Fraction of $CD94^+$NK cells among $Lin^-TBX21^+$ PBMCs from people who are HIV-$1^-$, n=13; HIV-$1^+$ with viremia >$10^4$/ml (HIV-$1^{high}$), n=8; or HIV-$1^+$ with viremia <$10^2$/ml (HIV-$1^{low}$), n=9.

(E) Fraction of $CD94^+$NK cells among $Lin^-TBX21^+$ PBMCs from individuals who are HIV-$1^-$, n=13; HIV-$1^+$ with $CD4^+$T cells >500/$mm^3$, n=7; or HIV-$1^+$ with $CD4^+$T cells <500/$mm^3$, n=10.

(F) Fraction of $CD94^+$NK cells among $Lin^-TBX21^+$ PBMCs after stimulation with PMA and ionomycin (n=10) or with IL-15 (n=10), or with IL-12+IL15 (n=4).

(G) Sorting strategy for $CD94^-$ and $CD94^+$NK cells.

(H) Percent CD107a among $CD94^-$ and $CD94^+$NK cells after PBMCs were stimulated with PMA/iono (n=5).

(I) Percent specific lysis of K562 or Jurkat cells by sorted $CD94^-$NK cells and $CD94^+$NK cells (n=8).

(J) Percent Ki67 and Annexin V among $CD94^-$ or $CD94^+$NK cells after the indicated treatment (n=4).

(K) Flow cytometry for KIR2DL1 as detected in Figure 2H.

Data is mean ± s.e.m. Each dot represents a unique sample. (C), two-tailed unpaired *t*-test; (D and E), Mann Whitney test; (F, H, I, and J), two-tailed paired *t*-test, lines connect cells from common donor. ns, non-significant, $^*p<0.05$, $^{**}p<0.01$, $^{***}p<0.001$.

**Figure S3. Single Cell Analysis of $CD94^-$ and $CD94^+$ NK Cells. Related to Figure 3.**

(A) Heatmap of 1,729 $CD94^-$ (blue) and 1,548 $CD94^+$NK cells (yellow) collected from 2 donors using all differentially expressed genes based on CD94 positivity.

(B) Plot of predictive strength as a function of the number of clusters in Figure 3B shows that 2 clusters yields stable and significant groupings, while separation into additional clusters artificially segregates the cells. The predictive strength was calculated using spectral clustering on the ICA components.

(C) Heatmap from Figure 3C was reconstructed utilizing the pseudotime ordering of single cells based on the minimum spanning tree.

(D) Flow cytometry for CD44, CXCR3, SELL, and CXCR6 on $TCF7^-$ and $TCF7^+$ NK cells.

(E) Flow cytometry for GZMK after sorted $Lin^-CD56^+CD94^-$NK cells were treated as in Figure 3H.

(F, G) PBMCs were treated with or without IL-15 for 5 days, $Lin^-CD56^+$ cells were gated on CD56 and CD94 (F) and percent $CD56^{hi}$ NK cells (G) (n=10). Data is mean ± s.e.m. two-tailed paired *t*-test, $^{***}$p<0.001.

**Figure S4. Distinct Chromatin Landscape of $CD94^+CD56^{hi}$ NK Cells. Related to Figure 5.**

(A) PCA based on H3K4me3 CUT&RUN of indicated NK subsets.

(B) Correlation between differentially expressed genes and enriched H3K4me3 regions by CUT&RUN (log2 fold change). Linear regression was applied to each plot, $r^2$ was as indicated, p<0.0001 for both the correlation and for the slope being significantly non-zero.

(C) Differential signals for H3K4me3 CUT&RUN and ATAC-Seq at the indicated loci in the indicated NK cell subsets.

(D) Overlapping signal for TCF7 CUT&RUN and ATAC-Seq at the indicated loci.

**Figure S5. Memory-Associated Gene Loci are Accessible in the $CD94^+CD56^{hi}$ NK Cell Subset. Related to Figure 6.**

(A) H3K4me1 and H3K4me3 CUT&RUN and ATAC-Seq signal on genes associated with memory T and NK cells, except for effector marker KLRG1, on the indicated NK cell subsets.

(B) IFN-γ production among $CD56^{dim}$ and $CD56^{hi}$ NK cells after stimulation with IL-12+IL-15 for 16 hr (n=4). mean ± s.e.m.; two tailed paired t-test; $^{**}$p<0.01.

(C) H3K4me1 and H3K4me3 CUT&RUN and ATAC-Seq signal at loci for IFN-γ signaling related genes.

**Figure S6. Loci for Genes Associated with WNT Signaling are Open in the $CD94^+CD56^{hi}$ NK Cell Subset. Related to Figure 6.**

H3K4me3 CUT&RUN and ATAC-Seq signal for gene loci of WNT signaling components and WNT target genes in the indicated NK cell subsets. AXIN1, in contrast, is a WNT inhibitory gene.

**Figure S7. WNT Inhibition Blocks Cytokine induced NK Cell Memory. Related to Figure 7.**

(A) Lin$^-$CD56$^+$ cells were treated as in Figure 7A. IFN-γ production was detected after the indicated treatment.

(B) PBMCs were treated with or without LGK974 for 16 hrs. Percentage of Lin$^-$TBX21$^+$ cells, and the CD94$^-$ and CD94$^+$ cells among the Lin$^-$TBX21$^+$ population, are indicated (left); data are representative of 4 donors. PBMCs were stimulated with IL-12 and IL-15 for 16 hrs in the absence or presence of LGK974. Live cells and Lin$^-$CD56$^+$ cells were examined (right); data are representative of 10 donors.

(C) PBMCs were treated with IL-12 and IL-15, or without (Control), in the presence or absence of LGK974 or M110, as indicated, for 16 hrs. CD56 and CD94 among Lin$^-$CD56$^+$ cells were checked. Data are representative of 4 donors.

(D, E) PBMCs were treated as indicated and the percent IFN-γ$^+$ among Lin$^-$TBX21$^+$ cells is shown (D, n=8; E, n=13), each dot represents a unique donor.

(F, G) Lin$^-$CD56$^+$ cells were transduced with shRNA as in Figure 7H and TCF7 was detected by flow cytometry.

(H) NKG2C was detected on Lin$^-$TBX21$^+$ cells from HIV-1$^-$ and HIV-1$^+$ individuals.

Data is mean ± s.e.m; two tailed paired t-test; ns, not significant, $^{***}p<0.001$.

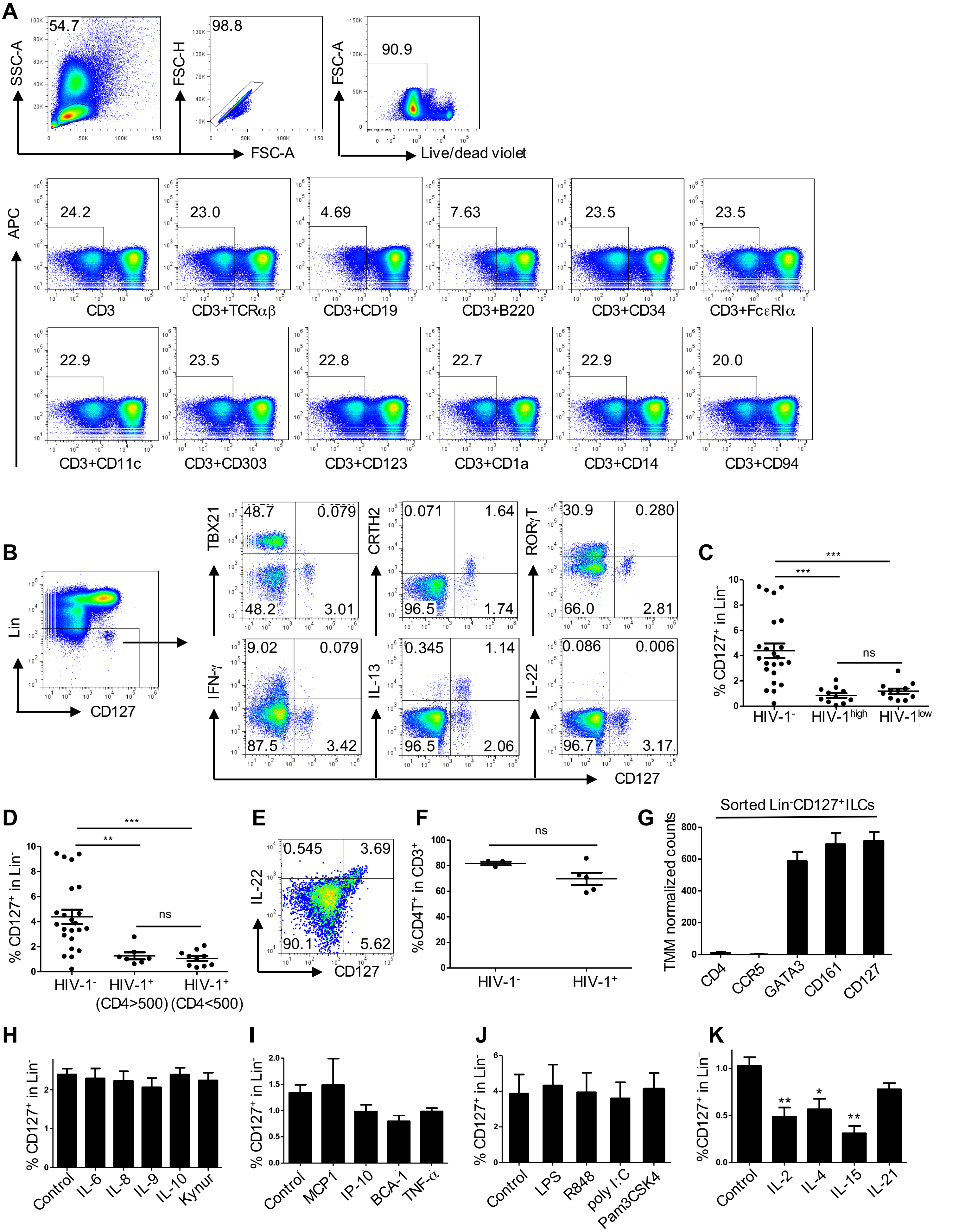

**Wang et al, Figure S1**

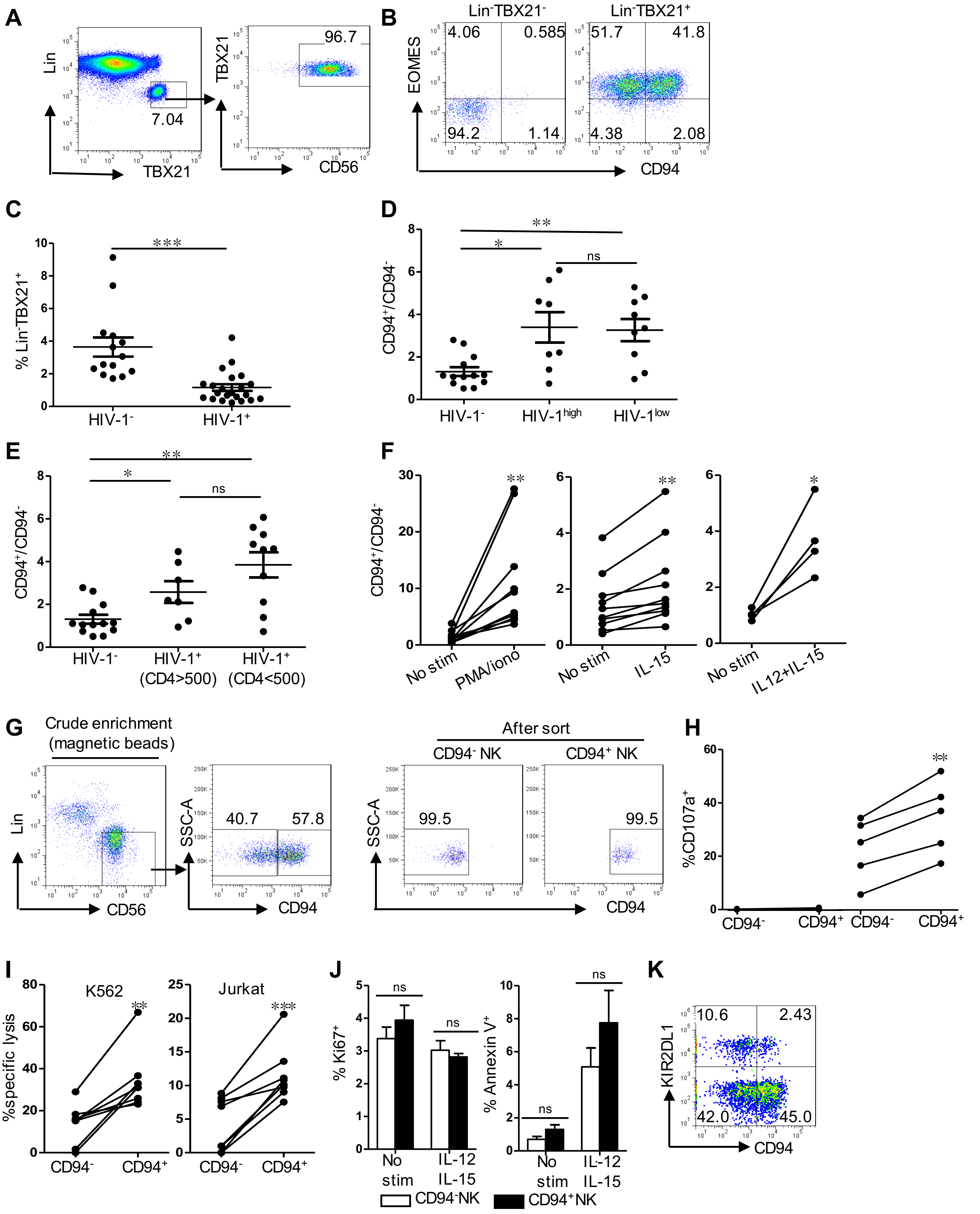

**Wang et al, Figure S2**

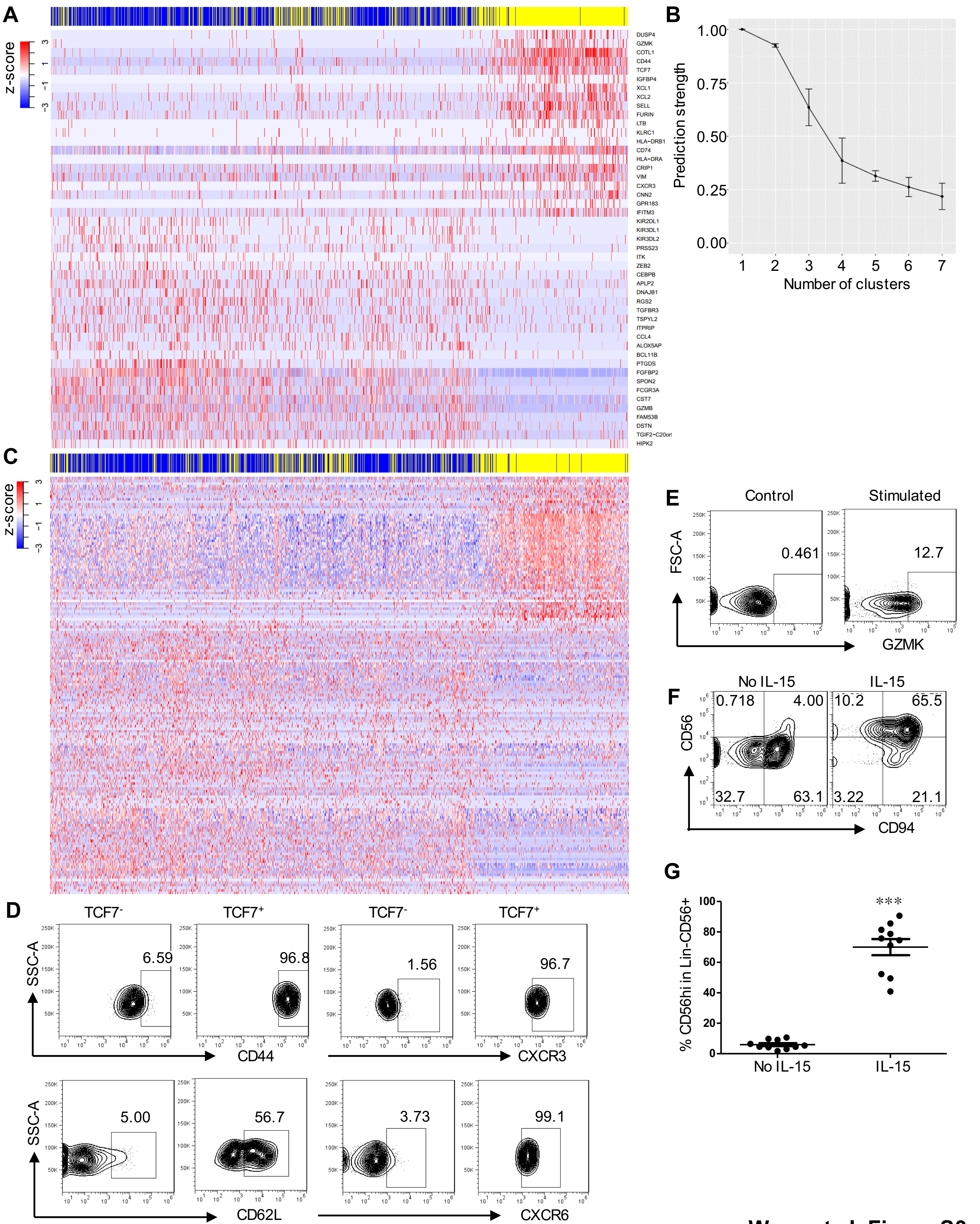

**Wang et al, Figure S3**

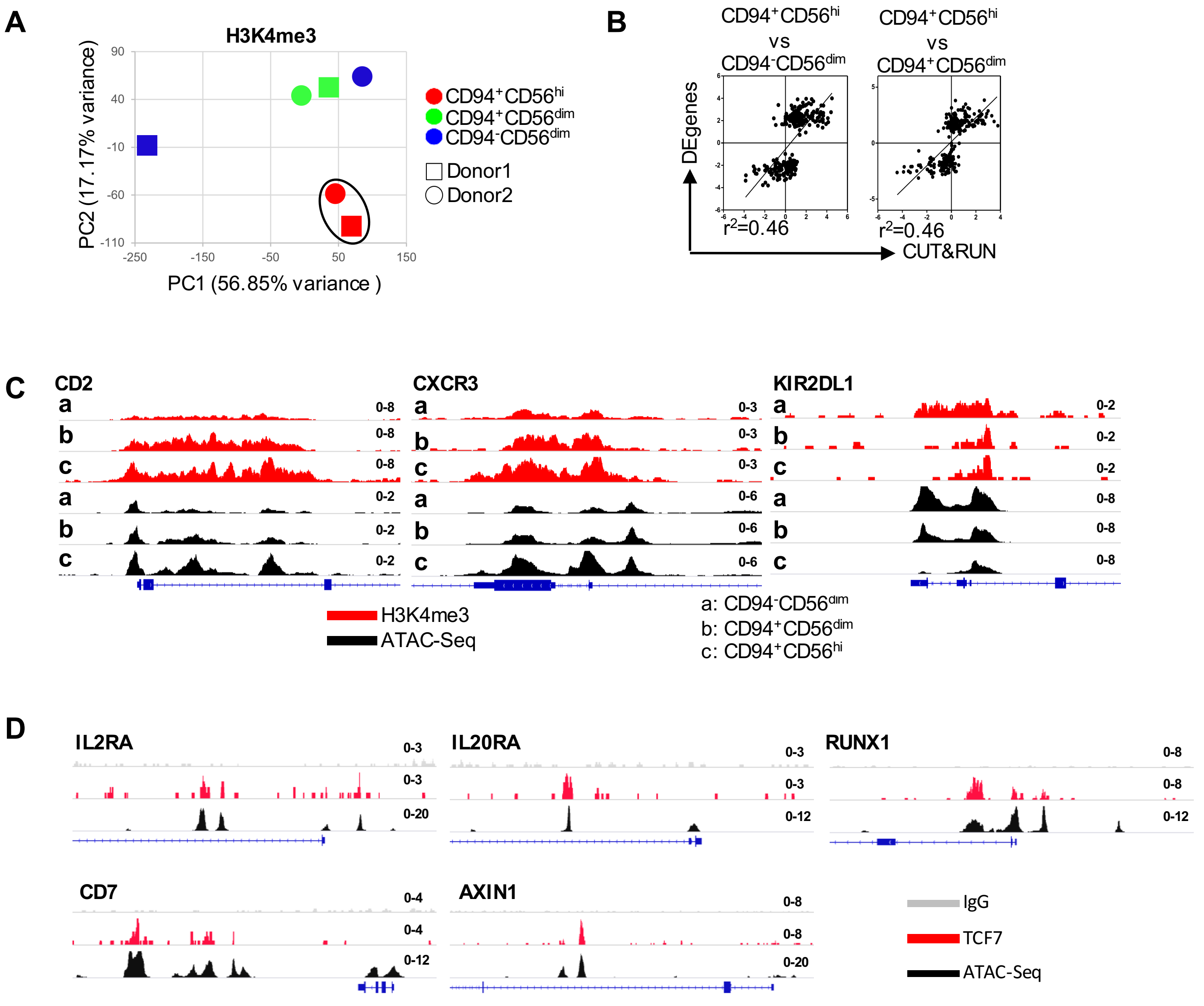

**Wang et al, Figure S4**

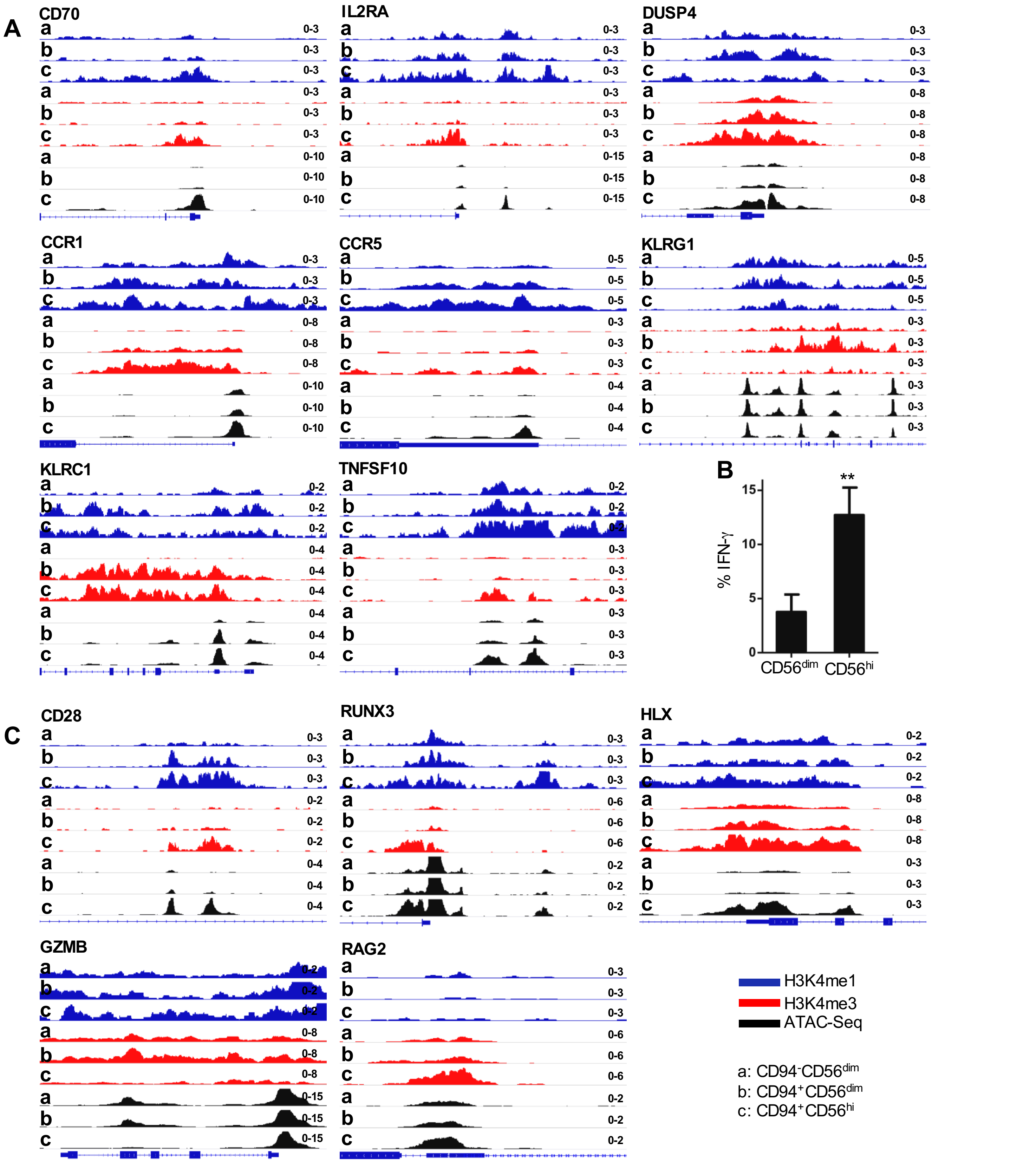

Wang et al, Figure S5

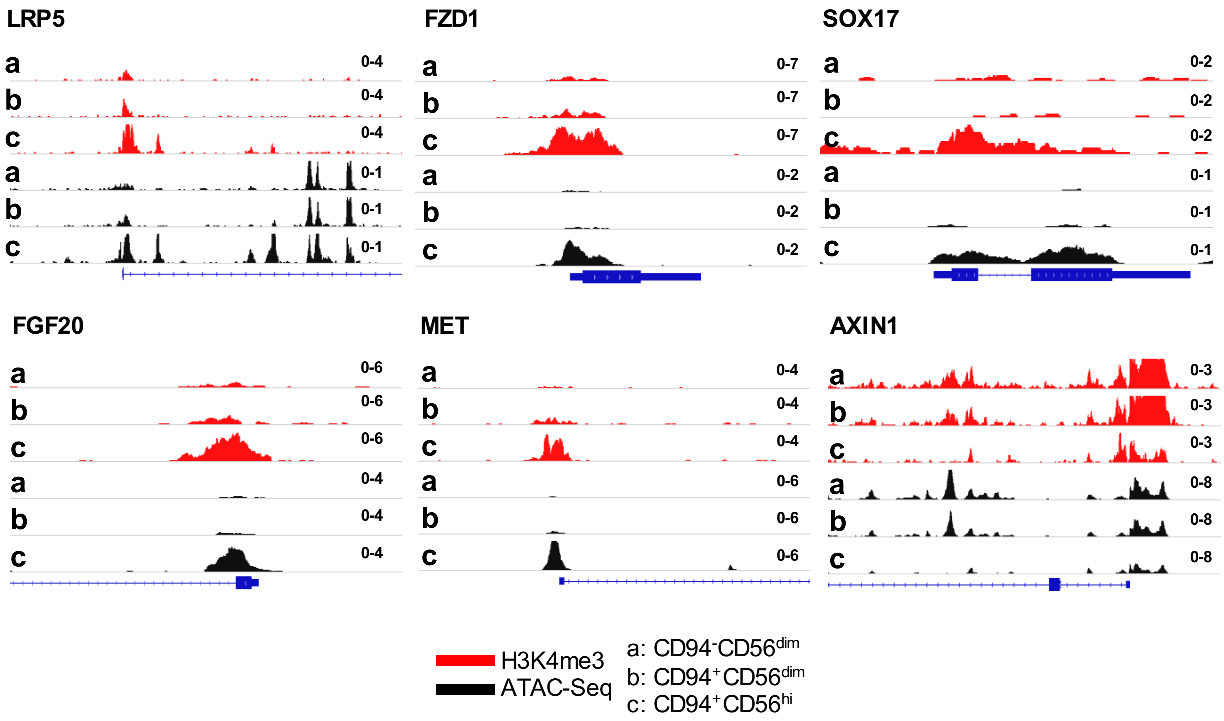

**Wang et al, Figure S6**

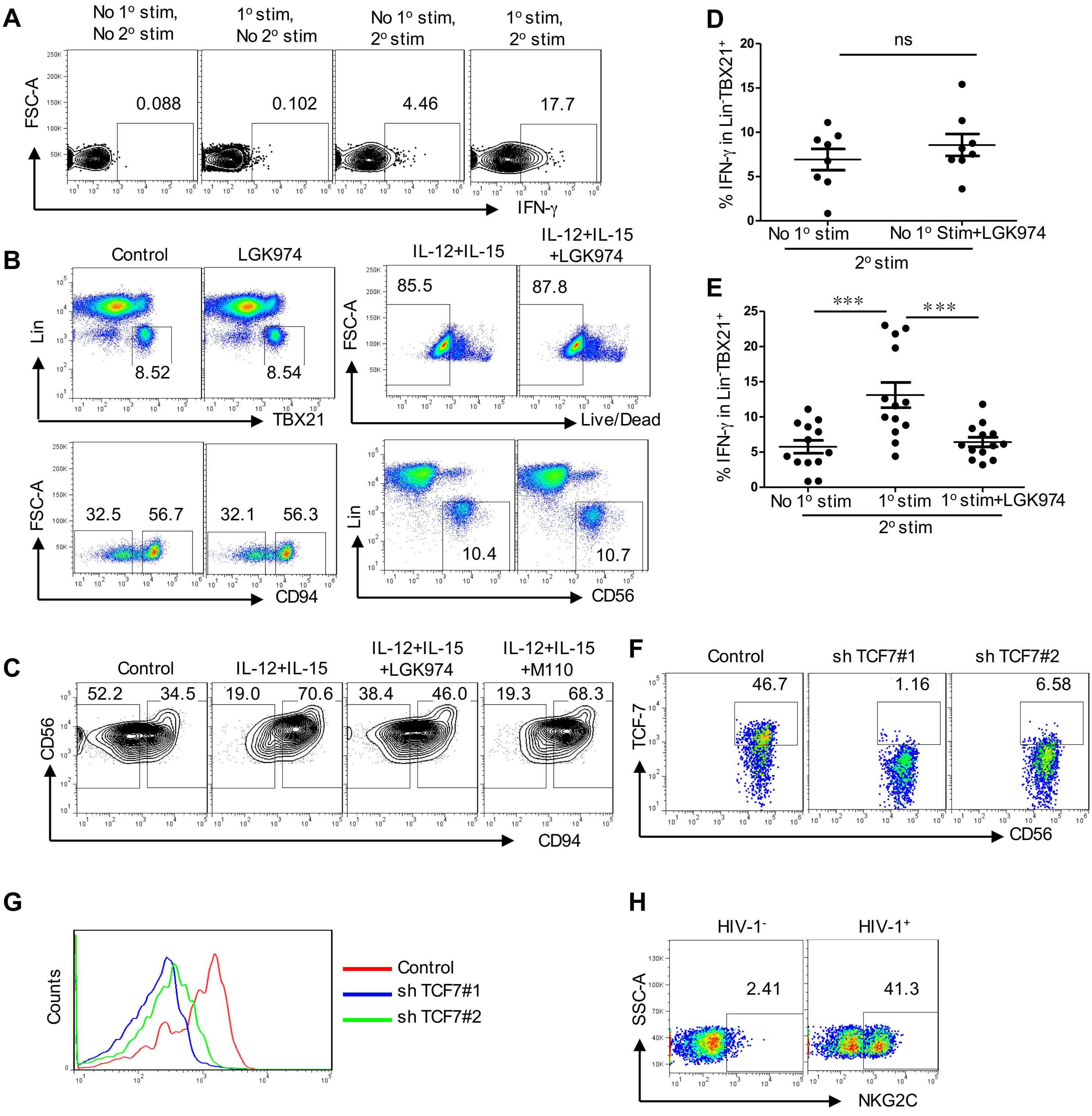

Wang et al, Figure S7

**Table S1 | Characteristics of HIV-1$^{+}$ donors**

| Sample ID# | Tissue source | CD4 (counts/mm$^3$) | Viral load (copies/ml) |
|---|---|---|---|
| 1 | colon/PBMC | 113 | < 20 |
| 2 | colon/PBMC | 620 | < 20 |
| 3 | colon/PBMC | 357 | 28 |
| 4 | colon/PBMC | 70 | < 20 |
| 5 | colon/PBMC | 787 | < 20 |
| 6 | colon/PBMC | 712 | < 20 |
| 7 | colon/PBMC | 764 | < 20 |
| 8-1 | PBMC | 246 | 73108 |
| 8-2 | PBMC | 256 | 578 |
| 9-1 | PBMC | 458 | 160070 |
| 9-2 | PBMC | N/A | 50 |
| 10-1 | PBMC | 282 | 12831 |
| 10-2 | PBMC | 567 | 55 |
| 11 | PBMC | 809 | 50 |
| 12-1 | PBMC | 436 | 373468 |
| 12-2 | PBMC | 1241 | 50 |
| 13-1 | PBMC | 281 | 18139 |
| 13-2 | PBMC | 425 | 50 |
| 14-1 | PBMC | 517 | 17525 |
| 14-2 | PBMC | 1108 | 50 |
| 15-1 | PBMC | 596 | 33833 |
| 15-2 | PBMC | 766 | 47748 |
| 15-3 | PBMC | 561 | 52015 |
| 15-4 | PBMC | 753 | 297 |
| 15-5 | PBMC | 649 | 50 |
| 15-6 | PBMC | 714 | 835 |
| 16-1 | PBMC | 161 | 94648 |
| 16-2 | PBMC | 622 | 57 |
| 17-1 | PBMC | 259 | 64081 |
| 17-2 | PBMC | 373 | 50 |
| 18-1 | PBMC | 13 | 484018 |
| 18-2 | PBMC | 51 | 50 |
| 19-1 | PBMC | 238 | 10494 |
| 19-2 | PBMC | 406 | 50 |
| 20-1 | PBMC | N/A | $>10^4$ |
| 20-2 | PBMC | N/A | $<10^3$ |
| 21-1 | PBMC | N/A | $>10^5$ |
| 22-1 | PBMC | N/A | 358588 |
| 22-2 | PBMC | 34 | 150 |
| 23-1 | PBMC | N/A | 29500 |
| 23-2 | PBMC | 322 | 50 |
| 24-1 | PBMC | 377 | 1264 |
| 24-2 | PBMC | 509 | 578 |
| 25 | PBMC | 474 | 50 |
| 26 | PBMC | 408 | 50 |
| 27 | PBMC | 308 | 50 |
| 28 | PBMC | 683 | 50 |

| | | | |
|---|---|---|---|
| 29 | PBMC | 313 | 50 |
| 30 | PBMC | 221 | 50 |
| 31 | PBMC | 772 | 50 |
| 32 | PBMC | 656 | 50 |
| 33 | PBMC | 732 | 50 |
| 34 | PBMC | 341 | 50 |
| N/A: data not available | | | |

| Table S2 \| DE genes from bulk RNA-Seq ($CD94^-$ VS $CD94^+$) | | | |
|---|---|---|---|
| **Gene** | **log2FoldChange** | **pvalue** | **padj** |
| KLRC1 | -4.15 | 3.47E-36 | 3.78E-32 |
| HOXA3 | -3.37 | 1.20E-21 | 4.36E-18 |
| RUNX2 | -3.27 | 2.50E-20 | 6.81E-17 |
| XCL1 | -2.94 | 2.15E-32 | 1.17E-28 |
| DUSP4 | -2.82 | 5.51E-20 | 1.20E-16 |
| HOXA9 | -2.62 | 3.63E-13 | 3.94E-10 |
| PACSIN1 | -2.57 | 1.04E-12 | 9.38E-10 |
| GPR183 | -2.51 | 1.71E-13 | 2.07E-10 |
| FAM167A | -2.48 | 6.37E-12 | 5.01E-09 |
| KLRC2 | -2.48 | 1.21E-13 | 1.65E-10 |
| FLNB | -2.34 | 1.09E-11 | 7.87E-09 |
| CABLES1 | -2.33 | 1.70E-10 | 8.04E-08 |
| TTN | -2.28 | 4.13E-10 | 1.87E-07 |
| ANO9 | -2.25 | 6.84E-10 | 2.75E-07 |
| TP53I11 | -2.20 | 1.42E-09 | 5.33E-07 |
| SCML1 | -2.11 | 2.75E-11 | 1.66E-08 |
| ATP1B1 | -2.08 | 1.51E-10 | 7.57E-08 |
| AGPAT5 | -2.07 | 9.52E-11 | 5.36E-08 |
| DLL1 | -2.06 | 1.53E-10 | 7.57E-08 |
| SPINT2 | -2.04 | 1.67E-08 | 5.34E-06 |
| PPP1R9A | -2.03 | 5.86E-10 | 2.45E-07 |
| HOXA10 | -1.99 | 2.53E-08 | 7.85E-06 |
| MICALCL | -1.94 | 4.87E-13 | 4.82E-10 |
| TCF7 | -1.90 | 1.68E-11 | 1.08E-08 |
| FXYD2 | -1.90 | 1.87E-07 | 5.36E-05 |
| FXYD6-FXYD2 | -1.90 | 1.87E-07 | 5.36E-05 |
| CDHR1 | -1.84 | 3.20E-07 | 7.90E-05 |
| NOD2 | -1.79 | 3.79E-07 | 9.16E-05 |
| RPS6KA2 | -1.78 | 8.49E-07 | 0.000177585 |
| LTA | -1.77 | 1.14E-06 | 0.000219504 |
| HVCN1 | -1.73 | 1.60E-11 | 1.08E-08 |
| SPIN3 | -1.70 | 2.58E-06 | 0.000418476 |
| TIE1 | -1.70 | 3.30E-06 | 0.000521978 |
| SIRPG | -1.69 | 1.94E-06 | 0.00033525 |
| CAPG | -1.69 | 2.28E-06 | 0.000381263 |
| SV2A | -1.68 | 4.03E-06 | 0.000576623 |
| COL9A2 | -1.67 | 3.60E-06 | 0.000538407 |
| NFIX | -1.65 | 5.73E-06 | 0.000763386 |
| FOXC1 | -1.64 | 3.99E-06 | 0.000576623 |
| COTL1 | -1.62 | 8.31E-06 | 0.001026496 |
| IER3 | -1.60 | 3.76E-09 | 1.32E-06 |
| SPTSSB | -1.60 | 6.10E-06 | 0.000798988 |
| TMEM154 | -1.59 | 7.83E-06 | 0.000977991 |

| | | | |
|---|---|---|---|
| CD44 | -1.59 | 6.45E-12 | 5.01E-09 |
| SELL | -1.57 | 1.63E-05 | 0.00184986 |
| DFNB31 | -1.56 | 5.76E-06 | 0.000763386 |
| CD83 | -1.56 | 9.87E-11 | 5.36E-08 |
| NELL2 | -1.55 | 1.76E-05 | 0.001975663 |
| NBL1 | -1.55 | 2.35E-07 | 6.55E-05 |
| TCF4 | -1.55 | 1.55E-06 | 0.000277084 |
| DTX1 | -1.53 | 2.27E-05 | 0.002443383 |
| HAPLN3 | -1.51 | 1.36E-05 | 0.001567283 |
| LGALS3 | -1.49 | 6.18E-06 | 0.000799295 |
| KIAA0754 | -1.45 | 4.11E-05 | 0.003904399 |
| COL4A3 | -1.45 | 3.46E-05 | 0.003479013 |
| FZD6 | -1.44 | 5.58E-05 | 0.005012489 |
| MSRB3 | -1.44 | 7.04E-05 | 0.006071759 |
| FUT8 | -1.42 | 5.01E-05 | 0.004619111 |
| GNA15 | -1.42 | 9.11E-05 | 0.00750193 |
| PTK2 | -1.41 | 0.000107871 | 0.00862253 |
| LMNA | -1.41 | 8.17E-09 | 2.78E-06 |
| MINOS1-NBL1 | -1.41 | 1.16E-06 | 0.000219504 |
| RFX2 | -1.40 | 6.73E-05 | 0.005850859 |
| RDH13 | -1.40 | 7.78E-05 | 0.006553685 |
| PLA2G6 | -1.38 | 1.11E-06 | 0.000219504 |
| MAML3 | -1.36 | 0.000175224 | 0.013320729 |
| CXCR3 | -1.35 | 2.51E-07 | 6.83E-05 |
| AGK | -1.34 | 2.24E-05 | 0.002443383 |
| AMPD3 | -1.32 | 0.000199959 | 0.01499141 |
| MCRS1 | -1.32 | 3.75E-05 | 0.003664616 |
| FHL1 | -1.30 | 0.000293258 | 0.018976261 |
| ACSL6 | -1.30 | 0.000363193 | 0.022806313 |
| SLC7A1 | -1.28 | 0.000437853 | 0.026235228 |
| USP35 | -1.28 | 0.000194462 | 0.014680521 |
| TIMP2 | -1.28 | 0.000442776 | 0.026290229 |
| CELSR1 | -1.28 | 0.000225721 | 0.016250427 |
| CD2 | -1.27 | 4.59E-06 | 0.000648114 |
| TTC7A | -1.26 | 0.000257563 | 0.017721294 |
| CLDND1 | -1.25 | 8.72E-16 | 1.58E-12 |
| CNTD1 | -1.24 | 0.00041543 | 0.025467584 |
| TOX2 | -1.24 | 0.000674705 | 0.035657857 |
| FAM102A | -1.23 | 6.67E-06 | 0.000853425 |
| CERKL | -1.21 | 0.000517222 | 0.029284981 |
| STRBP | -1.20 | 0.000228657 | 0.016353464 |
| CDK6 | -1.20 | 3.58E-05 | 0.003534833 |
| ITM2C | -1.19 | 0.000808857 | 0.041282097 |
| CXXC5 | -1.19 | 4.50E-10 | 1.96E-07 |
| GZMK | -1.18 | 0.00056434 | 0.031174354 |

| | | | |
|---|---|---|---|
| MAN1C1 | -1.18 | 0.000703939 | 0.03661492 |
| YBX3 | -1.16 | 0.001401409 | 0.061183583 |
| ARHGAP10 | -1.16 | 0.001366771 | 0.060626524 |
| GAS7 | -1.16 | 7.67E-05 | 0.006516999 |
| ANKRD27 | -1.16 | 0.000126013 | 0.009855332 |
| GABPB1 | -1.14 | 0.000272468 | 0.018397546 |
| IL18R1 | -1.14 | 5.63E-07 | 0.00012489 |
| TSPO | -1.14 | 8.93E-06 | 0.001066794 |
| BACH2 | -1.14 | 0.000523562 | 0.029490361 |
| CCR7 | -1.14 | 0.001718584 | 0.069711675 |
| HLA-DMA | -1.13 | 0.000325294 | 0.020924693 |
| CNN2 | -1.12 | 0.000141787 | 0.011009729 |
| MICAL2 | -1.12 | 0.000444982 | 0.026290229 |
| KLHL23 | -1.12 | 0.00158146 | 0.066123289 |
| NAT9 | -1.12 | 0.001362081 | 0.060626524 |
| DPH2 | -1.11 | 0.002330978 | 0.087986327 |
| UBL7 | -1.11 | 0.001332604 | 0.05986256 |
| GNAQ | -1.10 | 0.000638699 | 0.034372769 |
| SLC22A15 | -1.10 | 0.001294262 | 0.058869978 |
| PDE6G | -1.10 | 0.002036639 | 0.078511706 |
| AP1S2 | -1.10 | 0.001511173 | 0.063982424 |
| SYPL1 | -1.10 | 0.000347826 | 0.021983799 |
| NIPSNAP3A | -1.09 | 0.002527454 | 0.092194521 |
| IL4R | -1.09 | 1.91E-05 | 0.002119167 |
| HLA-DMB | -1.09 | 0.002641049 | 0.094755268 |
| DPM2 | -1.08 | 0.002724092 | 0.095861625 |
| GADD45G | -1.07 | 0.003006834 | 0.103114493 |
| IGFBP4 | -1.07 | 0.00161151 | 0.066611105 |
| E2F3 | -1.07 | 0.001325863 | 0.059806889 |
| AHR | -1.07 | 0.001383072 | 0.060626524 |
| SGSM2 | -1.06 | 5.80E-05 | 0.005129973 |
| TYMS | -1.06 | 0.001758345 | 0.070275621 |
| MEX3B | -1.06 | 0.002827187 | 0.098557982 |
| TLE3 | -1.05 | 5.43E-06 | 0.000737896 |
| ZNF296 | -1.05 | 0.000942542 | 0.045539445 |
| GAB1 | -1.05 | 0.003389775 | 0.111330048 |
| RASSF2 | -1.05 | 0.004157052 | 0.127407939 |
| C2CD2 | -1.04 | 0.003261103 | 0.108746778 |
| HLA-DRB1 | -1.04 | 0.000895881 | 0.044724015 |
| GNPDA1 | -1.04 | 0.004268513 | 0.129617345 |
| C4orf32 | -1.04 | 0.001774311 | 0.070653977 |
| ZNF8 | -1.03 | 0.002463261 | 0.090896601 |
| PLP2 | -1.03 | 1.32E-08 | 4.35E-06 |
| PHACTR2 | -1.03 | 4.41E-07 | 0.000103247 |
| ADCY3 | -1.03 | 0.004725058 | 0.137710744 |

| | | | |
|---|---|---|---|
| KDSR | -1.02 | 0.000283885 | 0.018817791 |
| ANKRD6 | -1.02 | 0.005058462 | 0.143730703 |
| SPATS2L | -1.02 | 0.005063827 | 0.143730703 |
| INPP4B | -1.02 | 0.002657278 | 0.095023926 |
| LRRC8D | -1.01 | 0.004160594 | 0.127407939 |
| HOOK1 | -1.01 | 0.005789202 | 0.156553278 |
| REC114 | -1.00 | 0.005285418 | 0.147327639 |
| TBX19 | 1.01 | 0.005828044 | 0.156823445 |
| NR1D1 | 1.01 | 0.004761999 | 0.138416287 |
| ARFIP1 | 1.01 | 0.005163576 | 0.144673287 |
| ERBB2 | 1.01 | 0.004710389 | 0.137710744 |
| TSPYL5 | 1.02 | 0.004690806 | 0.137710744 |
| GCLM | 1.02 | 4.06E-05 | 0.003901578 |
| GMNN | 1.03 | 0.002997945 | 0.103114493 |
| PDXP | 1.03 | 0.000614372 | 0.033560761 |
| ARL5B | 1.03 | 0.004663288 | 0.137383742 |
| MGAM | 1.03 | 0.004646685 | 0.137383742 |
| PPP2CB | 1.03 | 0.001921133 | 0.075395793 |
| CCL4 | 1.03 | 9.04E-05 | 0.00750193 |
| CTNNA1 | 1.03 | 0.004501358 | 0.134434802 |
| SMG5 | 1.04 | 0.001690268 | 0.069058597 |
| SORBS3 | 1.04 | 0.003943078 | 0.123129857 |
| ABI3 | 1.05 | 0.000291158 | 0.018953163 |
| HECTD3 | 1.05 | 0.000564929 | 0.031174354 |
| KRT73 | 1.05 | 0.003602412 | 0.116553024 |
| IL1RAP | 1.06 | 0.003498428 | 0.113866491 |
| EML6 | 1.07 | 0.001754952 | 0.070275621 |
| KIFC3 | 1.07 | 0.000611831 | 0.033560761 |
| RUNDC1 | 1.07 | 0.000384167 | 0.023864469 |
| FGF9 | 1.07 | 0.00281316 | 0.098557982 |
| ZNF251 | 1.09 | 0.002718852 | 0.095861625 |
| PLOD1 | 1.09 | 0.000621462 | 0.033611518 |
| CCT6B | 1.09 | 0.002721038 | 0.095861625 |
| FGFBP2 | 1.10 | 3.31E-06 | 0.000521978 |
| NME8 | 1.11 | 0.000439225 | 0.026235228 |
| OTUD7B | 1.11 | 0.001696131 | 0.069058597 |
| DBNDD2 | 1.11 | 2.34E-06 | 0.000385029 |
| ODF2 | 1.12 | 0.000151052 | 0.011563979 |
| MBD4 | 1.13 | 4.91E-07 | 0.000111134 |
| TCEANC | 1.13 | 0.001178943 | 0.054178865 |
| FBXL14 | 1.13 | 0.001383002 | 0.060626524 |
| MTSS1 | 1.14 | 3.20E-05 | 0.003312325 |
| PPM1L | 1.15 | 0.001618305 | 0.06663862 |
| XRCC6BP1 | 1.15 | 0.001154508 | 0.053635281 |
| CTSF | 1.16 | 2.87E-07 | 7.25E-05 |

| | | | |
|---|---|---|---|
| POU2F2 | 1.16 | 1.17E-06 | 0.000219504 |
| RBPMS2 | 1.17 | 0.00118116 | 0.054178865 |
| WDR89 | 1.18 | 0.001125168 | 0.052496573 |
| ZNF441 | 1.19 | 0.000675699 | 0.035657857 |
| LGR6 | 1.20 | 0.000907284 | 0.04483218 |
| CD6 | 1.20 | 3.78E-05 | 0.003664616 |
| PLEKHB1 | 1.22 | 0.000642498 | 0.034406866 |
| KIR2DL3 | 1.22 | 9.73E-05 | 0.007949306 |
| ZNF615 | 1.23 | 1.27E-06 | 0.000234302 |
| CAPN5 | 1.24 | 0.000698101 | 0.036485849 |
| SEPT8 | 1.24 | 0.000552235 | 0.030945109 |
| MARCKSL1 | 1.26 | 6.63E-05 | 0.005810012 |
| CYBRD1 | 1.27 | 0.000230204 | 0.016356486 |
| ZBTB8OS | 1.27 | 2.65E-07 | 7.01E-05 |
| KIR2DS4 | 1.32 | 7.59E-05 | 0.006493508 |
| MAN1A1 | 1.33 | 3.63E-06 | 0.000538407 |
| FCRL6 | 1.33 | 2.45E-15 | 3.81E-12 |
| GGACT | 1.35 | 0.000211623 | 0.015650007 |
| KIR2DL1 | 1.36 | 2.75E-05 | 0.002899312 |
| C16orf91 | 1.40 | 3.52E-05 | 0.003509004 |
| EBF4 | 1.40 | 0.000125903 | 0.009855332 |
| RASSF4 | 1.43 | 3.46E-06 | 0.000529424 |
| RNMTL1 | 1.46 | 1.80E-06 | 0.000316046 |
| LDLR | 1.46 | 1.52E-06 | 0.000274948 |
| INTS2 | 1.49 | 4.47E-05 | 0.00419081 |
| C12orf76 | 1.50 | 2.84E-05 | 0.002973602 |
| GAS1 | 1.53 | 9.71E-06 | 0.001147404 |
| DAB2 | 1.54 | 4.46E-07 | 0.000103247 |
| SPRED2 | 1.57 | 8.46E-06 | 0.001033237 |
| PRSS57 | 1.57 | 3.66E-06 | 0.000538407 |
| KRT72 | 1.74 | 2.81E-07 | 7.25E-05 |
| TK2 | 1.82 | 2.39E-09 | 8.66E-07 |
| 2-fold difference as a cut-off, p＜0.05 | | | |

| **Table S3. Single cell DEgenes based on CD94.** | | | |
|---|---|---|---|
| **Gene** | **logFC** | **PValue** | **FDR** |
| COTL1 | -2.194 | 1.60E-90 | 5.39E-87 |
| SELL | -1.869 | 2.66E-59 | 2.99E-56 |
| GZMK | -1.777 | 3.42E-84 | 6.92E-81 |
| CD44 | -1.524 | 3.33E-63 | 4.82E-60 |
| TCF7 | -1.491 | 8.42E-65 | 1.42E-61 |
| FURIN | -1.212 | 1.31E-36 | 6.29E-34 |
| XCL2 | -1.151 | 1.99E-35 | 9.15E-33 |
| XCL1 | -1.119 | 9.26E-43 | 6.25E-40 |
| CXCR3 | -1.055 | 1.86E-15 | 2.30E-13 |
| DUSP4 | -1.040 | 2.49E-37 | 1.33E-34 |
| FCGR3A | 1.056 | 1.10E-27 | 3.29E-25 |
| CCL4 | 1.223 | 5.00E-15 | 5.96E-13 |
| SPON2 | 1.304 | 1.84E-40 | 1.16E-37 |
| FGFBP2 | 1.702 | 1.70E-119 | 1.72E-115 |
| PTGDS | 1.836 | 1.33E-38 | 7.47E-36 |
| 2-fold difference as a cut-off, $p < 0.05$ | | | |

| Table S4 \| Single cell DEgenes based on spectral clustering | | | |
|---|---|---|---|
| **Gene** | **Log2(Fold Change)** | **pval** | **qval** |
| FGFBP2 | -4.12 | 1.93E-156 | 1.30E-153 |
| COTL1 | 3.99 | 3.78E-130 | 1.28E-127 |
| TCF7 | 3.67 | 2.56E-120 | 5.76E-118 |
| GZMK | 4.77 | 1.04E-89 | 1.76E-87 |
| CD44 | 2.51 | 6.37E-69 | 8.60E-67 |
| SELL | 2.45 | 1.62E-60 | 1.82E-58 |
| SPON2 | -4.33 | 1.92E-58 | 1.85E-56 |
| B2M | -0.6 | 7.21E-58 | 6.08E-56 |
| EEF1A1 | 0.78 | 5.28E-57 | 3.96E-55 |
| XCL2 | 2.22 | 4.80E-55 | 3.24E-53 |
| CD247 | -1.93 | 9.16E-49 | 5.15E-47 |
| RPS18 | 1.15 | 3.64E-47 | 1.89E-45 |
| IFITM3 | 2.38 | 3.66E-45 | 1.77E-43 |
| RPLP0 | 1.22 | 1.69E-42 | 7.14E-41 |
| CD74 | 1.59 | 3.06E-42 | 1.21E-40 |
| TPT1 | 0.89 | 1.39E-41 | 5.23E-40 |
| RPL10A | 1.04 | 4.09E-39 | 1.31E-37 |
| RPSA | 1.18 | 4.67E-39 | 1.43E-37 |
| RPS12 | 0.94 | 1.11E-37 | 3.26E-36 |
| CST7 | -2.06 | 2.94E-37 | 8.25E-36 |
| SYNE2 | -3.13 | 4.96E-36 | 1.34E-34 |
| FAM53B | -3.1 | 6.69E-36 | 1.74E-34 |
| RPS24 | 0.89 | 9.82E-36 | 2.46E-34 |
| PIM3 | 1.75 | 1.03E-35 | 2.47E-34 |
| EEF1G | 1.02 | 1.20E-35 | 2.80E-34 |
| RPL3 | 0.88 | 1.92E-35 | 4.33E-34 |
| GZMB | -2.13 | 5.89E-35 | 1.28E-33 |
| RPLP1 | 0.83 | 6.43E-34 | 1.32E-32 |
| NKG7 | -1.09 | 1.12E-33 | 2.21E-32 |
| FCGR3A | -2.97 | 1.28E-33 | 2.47E-32 |
| RPL13 | 0.77 | 1.77E-33 | 3.31E-32 |
| GNG2 | -2.77 | 2.81E-33 | 5.14E-32 |
| RPS3A | 0.86 | 3.01E-33 | 5.35E-32 |
| RPS6 | 0.95 | 1.05E-32 | 1.82E-31 |
| RPL13A | 0.72 | 7.29E-32 | 1.17E-30 |
| FURIN | 1.9 | 2.31E-31 | 3.54E-30 |
| RPL36A-HNRNPH2 | 1.12 | 5.39E-31 | 7.74E-30 |
| RPS8 | 0.99 | 1.05E-30 | 1.48E-29 |
| RPL31 | 0.82 | 1.87E-30 | 2.58E-29 |
| RPS5 | 0.86 | 4.16E-29 | 5.51E-28 |
| RPL36A | 0.92 | 2.14E-27 | 2.68E-26 |
| CCL5 | -1.41 | 2.15E-27 | 2.68E-26 |
| PRSS23 | -3.95 | 2.52E-26 | 2.99E-25 |
| HLA-B | -0.46 | 2.66E-26 | 3.10E-25 |
| TAGLN2 | 1.36 | 5.36E-26 | 6.13E-25 |
| RPL35 | 0.98 | 9.88E-26 | 1.11E-24 |
| FXYD5 | 1.36 | 1.04E-25 | 1.13E-24 |

| | | | |
|---|---|---|---|
| EEF1B2 | 0.89 | 6.72E-25 | 6.98E-24 |
| RPS9 | 0.71 | 2.69E-24 | 2.71E-23 |
| CRIP1 | 1.48 | 2.83E-24 | 2.81E-23 |
| CMC1 | 1.39 | 6.54E-24 | 6.21E-23 |
| RPL37 | 0.66 | 1.28E-23 | 1.20E-22 |
| HLA-C | -0.53 | 1.70E-23 | 1.55E-22 |
| ADGRG1 | -2.57 | 1.84E-23 | 1.66E-22 |
| RPS28 | 0.6 | 2.25E-23 | 2.00E-22 |
| FKBP11 | -2.84 | 1.34E-21 | 1.09E-20 |
| S100A4 | -1.64 | 3.49E-21 | 2.77E-20 |
| S100A6 | -1.46 | 7.21E-21 | 5.53E-20 |
| CEBPB | -2.49 | 1.44E-20 | 1.06E-19 |
| RPL12 | 0.76 | 4.60E-20 | 3.20E-19 |
| ARPC2 | -1.19 | 1.72E-19 | 1.13E-18 |
| RGS2 | -3.16 | 1.82E-19 | 1.18E-18 |
| ITGB2 | -1.15 | 2.64E-19 | 1.70E-18 |
| APLP2 | -2.71 | 5.43E-19 | 3.46E-18 |
| GZMH | -2.28 | 7.22E-19 | 4.47E-18 |
| ARL4C | -0.99 | 6.18E-18 | 3.66E-17 |
| RPL10 | 0.42 | 6.25E-18 | 3.67E-17 |
| HLA-E | -0.71 | 9.45E-18 | 5.45E-17 |
| GZMA | -1.59 | 1.45E-17 | 8.25E-17 |
| GATA3 | 0.8 | 2.46E-17 | 1.36E-16 |
| DSTN | -2.23 | 2.64E-17 | 1.45E-16 |
| GNPTAB | -2.24 | 1.59E-16 | 8.57E-16 |
| RPS16 | 0.49 | 7.78E-16 | 3.98E-15 |
| RGCC | 1.32 | 1.89E-15 | 9.47E-15 |
| EFHD2 | -1.02 | 4.35E-15 | 2.16E-14 |
| IGFBP7 | -2.23 | 1.05E-14 | 5.05E-14 |
| PRF1 | -1.85 | 6.36E-14 | 2.94E-13 |
| HSPA5 | -1.22 | 1.41E-13 | 6.45E-13 |
| ALOX5AP | -2.98 | 4.19E-13 | 1.86E-12 |
| EIF3E | 1 | 4.54E-13 | 2.00E-12 |
| MTRNR2L10 | 0.62 | 7.74E-13 | 3.37E-12 |
| ABHD17A | -1.19 | 1.37E-12 | 5.80E-12 |
| FLNA | -1.07 | 2.38E-12 | 9.81E-12 |
| BTF3 | 0.44 | 2.55E-12 | 1.04E-11 |
| APMAP | -1.39 | 5.57E-12 | 2.24E-11 |
| IFITM1 | 0.49 | 1.20E-11 | 4.75E-11 |
| RORA | -1.91 | 2.27E-11 | 8.80E-11 |
| GZMM | -1.03 | 2.60E-11 | 1.00E-10 |
| UBB | -0.89 | 3.52E-11 | 1.33E-10 |
| CFL1 | -0.87 | 6.27E-11 | 2.37E-10 |
| LCP1 | -1.14 | 9.25E-11 | 3.47E-10 |
| CYBA | -0.71 | 5.85E-10 | 2.12E-09 |
| PTMA | -0.44 | 6.00E-10 | 2.16E-09 |
| NAP1L1 | 0.56 | 6.08E-10 | 2.18E-09 |
| CLK1 | -1.29 | 6.36E-10 | 2.26E-09 |
| RPS26 | 0.5 | 6.48E-10 | 2.28E-09 |

| | | | |
|---|---|---|---|
| GNLY | 0.29 | 8.52E-10 | 2.96E-09 |
| DNAJB1 | -1.79 | 8.70E-10 | 3.01E-09 |
| SLC15A4 | -1.65 | 1.04E-09 | 3.58E-09 |
| TGIF2-C20orf24 | -1.71 | 1.38E-09 | 4.69E-09 |
| CELF2 | -1.47 | 1.48E-09 | 5.00E-09 |
| VIM | 0.59 | 1.58E-09 | 5.30E-09 |
| ACTB | -0.79 | 1.62E-09 | 5.37E-09 |
| NR4A2 | -1.32 | 2.02E-09 | 6.68E-09 |
| TSPYL2 | -2.05 | 3.45E-09 | 1.13E-08 |
| MT2A | -1.3 | 3.66E-09 | 1.19E-08 |
| MTRNR2L1 | 0.24 | 8.87E-09 | 2.86E-08 |
| ATP1B3 | -1.27 | 1.84E-08 | 5.78E-08 |
| LYN | -1.65 | 2.65E-08 | 8.22E-08 |
| RAB8B | -1.12 | 3.93E-08 | 1.19E-07 |
| LRPAP1 | -0.37 | 7.92E-08 | 2.39E-07 |
| PSAP | -1.15 | 1.64E-07 | 4.79E-07 |
| EIF1 | -0.64 | 3.16E-07 | 8.85E-07 |
| GABARAPL1 | -1.19 | 5.46E-07 | 1.52E-06 |
| FCRL6 | -3.46 | 6.31E-07 | 1.75E-06 |
| TNFRSF1B | -0.86 | 1.15E-06 | 3.12E-06 |
| HLA-A | -0.4 | 1.34E-06 | 3.58E-06 |
| FTH1 | -0.59 | 1.38E-06 | 3.67E-06 |
| SORL1 | -0.87 | 7.73E-06 | 1.98E-05 |
| ARHGDIB | -0.88 | 8.78E-06 | 2.24E-05 |
| LAPTM5 | -0.64 | 1.20E-05 | 3.02E-05 |
| NFKBIA | 0.18 | 1.68E-05 | 4.19E-05 |
| SMAD7 | -0.06 | 2.05E-05 | 5.06E-05 |
| YWHAZ | -0.86 | 3.56E-05 | 8.62E-05 |
| CAPZB | -1.32 | 4.93E-05 | 1.18E-04 |
| SRSF5 | -0.82 | 5.08E-05 | 1.21E-04 |
| SLC7A5 | 0.15 | 6.47E-05 | 1.53E-04 |
| MGAT1 | -0.99 | 6.56E-05 | 1.55E-04 |
| IL2RG | -0.8 | 7.21E-05 | 1.68E-04 |
| PPP1CA | -0.37 | 1.20E-04 | 2.74E-04 |
| CALM1 | -0.63 | 1.38E-04 | 3.11E-04 |
| CD164 | -0.77 | 1.63E-04 | 3.62E-04 |
| CTSD | -0.66 | 1.76E-04 | 3.88E-04 |
| TPM3 | -0.71 | 2.33E-04 | 5.08E-04 |
| PDIA3 | -0.7 | 2.44E-04 | 5.27E-04 |
| DDX6 | -0.98 | 2.45E-04 | 5.29E-04 |
| HMGB1 | -0.79 | 2.75E-04 | 5.85E-04 |
| DBI | -0.95 | 3.89E-04 | 8.10E-04 |
| PTPRE | -1.44 | 5.11E-04 | 1.05E-03 |
| UBC | -0.53 | 5.77E-04 | 1.17E-03 |
| MXRA7 | -0.81 | 8.25E-04 | 1.62E-03 |
| TMA7 | -0.58 | 1.07E-03 | 2.08E-03 |
| CHD1 | -1.09 | 1.13E-03 | 2.19E-03 |
| PTPRCAP | -0.97 | 1.28E-03 | 2.46E-03 |
| CALR | -0.6 | 1.43E-03 | 2.73E-03 |

| | | | |
|---|---|---|---|
| KLRC3 | 0.6 | 1.71E-03 | 3.25E-03 |
| SH3BGRL3 | -0.71 | 1.82E-03 | 3.45E-03 |
| THUMPD1 | -0.99 | 2.48E-03 | 4.58E-03 |
| ZDHHC7 | 0.17 | 2.58E-03 | 4.75E-03 |
| RAB27A | -0.28 | 6.06E-03 | 1.05E-02 |
| KPNB1 | -0.65 | 6.71E-03 | 1.16E-02 |
| MYL6 | -0.71 | 7.32E-03 | 1.25E-02 |
| RNF216 | -1.11 | 8.40E-03 | 1.42E-02 |
| IFRD1 | 0.36 | 1.14E-02 | 1.88E-02 |
| UBALD2 | -0.51 | 1.51E-02 | 2.42E-02 |
| BCL9L | -0.57 | 1.62E-02 | 2.57E-02 |
| CXCR4 | -0.19 | 1.64E-02 | 2.59E-02 |
| RBM39 | -0.56 | 1.93E-02 | 3.00E-02 |
| ATPIF1 | -0.41 | 4.37E-02 | 6.24E-02 |
| CXCR3 | 3.37 | 6.23E-02 | 8.23E-02 |

| Table S5 \| 3 NK subsets RNA-Seq | | |
|---|---|---|
| DE genes from bulk RNA-Seq ($CD94^{+}CD56^{hi}$ VS $CD94^{-}CD56^{dim}$ ) | | |
| **Gene** | **log2FoldChange** | **padj** |
| IL7R | 3.994821053 | 5.92E-45 |
| SPTSSB | 3.958938484 | 9.38E-19 |
| TNFRSF11A | 3.920498682 | 7.89E-19 |
| FOXC1 | 3.908183536 | 1.06E-07 |
| ANO9 | 3.733987227 | 5.97E-14 |
| CDHR1 | 3.610342384 | 6.32E-10 |
| DUSP4 | 3.562557267 | 6.64E-11 |
| PPP1R9A | 3.464530615 | 3.77E-21 |
| HAPLN3 | 3.429462393 | 2.30E-08 |
| TSPAN18 | 3.397024978 | 1.44E-06 |
| NR4A3 | 3.36151622 | 6.37E-16 |
| GZMK | 3.36058433 | 4.30E-12 |
| SPINK2 | 3.343624452 | 2.24E-07 |
| KLRC1 | 3.342472979 | 7.78E-15 |
| TCF7 | 3.313149888 | 1.10E-30 |
| NFIX | 3.286516924 | 4.92E-08 |
| NELL2 | 3.26106256 | 7.66E-08 |
| PRDM8 | 3.228699383 | 8.66E-12 |
| GAB1 | 3.180728143 | 4.51E-10 |
| KIT | 3.080197859 | 2.87E-10 |
| SPRY2 | 3.070702405 | 1.08E-06 |
| DLL1 | 3.06144056 | 2.35E-13 |
| IL12RB2 | 3.040296792 | 4.07E-16 |
| RUNX2 | 2.95269978 | 1.21E-11 |
| PAWR | 2.879406515 | 5.80E-06 |
| DTX1 | 2.870887962 | 3.64E-05 |
| DENND5A | 2.815603157 | 2.64E-08 |
| GNAI1 | 2.811174017 | 0.000860713 |
| FUT7 | 2.795566255 | 5.36E-06 |
| XCL1 | 2.760973132 | 9.44E-08 |
| ADGRG3 | 2.753626887 | 7.75E-05 |
| PACSIN1 | 2.753145983 | 1.58E-07 |
| SIRPG | 2.751031504 | 1.39E-05 |
| CRTAM | 2.734483907 | 7.06E-08 |
| CDCP1 | 2.689577624 | 0.000263333 |
| COL9A2 | 2.688356198 | 2.14E-06 |
| HOXA9 | 2.669591342 | 0.000421496 |
| FLNB | 2.66158483 | 2.24E-07 |
| IGFBP4 | 2.657135057 | 4.82E-06 |
| TEC | 2.650337438 | 2.12E-05 |
| SSBP2 | 2.646412378 | 1.99E-08 |
| SLIT3 | 2.610678808 | 0.003426024 |

| | | |
|---|---|---|
| MMP25 | 2.591381748 | 0.000299707 |
| MYC | 2.590241329 | 9.61E-05 |
| LMNA | 2.564314609 | 0.000363815 |
| CAPG | 2.561015043 | 0.000102925 |
| CCR1 | 2.552903649 | 1.27E-05 |
| TNFSF11 | 2.524873444 | 0.000169449 |
| HVCN1 | 2.522387436 | 1.32E-10 |
| FXYD7 | 2.513886917 | 1.20E-05 |
| CASK | 2.494808061 | 3.35E-07 |
| TIE1 | 2.490144312 | 2.42E-06 |
| SPRY1 | 2.482902782 | 0.000402705 |
| LRRC75B | 2.481459618 | 0.000571666 |
| ZEB1 | 2.472857375 | 4.69E-09 |
| EPHA4 | 2.467706494 | 6.14E-09 |
| INPP4B | 2.452004165 | 0.000283845 |
| LEF1 | 2.441786385 | 2.59E-08 |
| EVA1B | 2.429775173 | 0.00044889 |
| HOXA5 | 2.427469432 | 0.001812047 |
| GRAMD1B | 2.42475269 | 0.001075106 |
| STYK1 | 2.409727125 | 3.04E-06 |
| SELL | 2.399798398 | 3.78E-06 |
| DACH1 | 2.384633943 | 0.002594235 |
| KLRC2 | 2.373964922 | 1.61E-06 |
| KIR2DL4 | 2.362047117 | 0.000177982 |
| TRIM47 | 2.359498491 | 0.002407911 |
| PI16 | 2.358012703 | 0.001277984 |
| CNR2 | 2.333605704 | 0.000894719 |
| CRACR2B | 2.319388962 | 0.001241282 |
| CXCR3 | 2.296086074 | 0.000106774 |
| CD44 | 2.265431018 | 2.35E-14 |
| DPP4 | 2.244531053 | 0.00113122 |
| HOXA10 | 2.238671457 | 0.001500488 |
| IER5L | 2.228554816 | 0.003261211 |
| MAML3 | 2.228510533 | 0.002206285 |
| BAIAP3 | 2.228136414 | 0.002552469 |
| RASSF8 | 2.227743078 | 0.00585985 |
| IL1RL1 | 2.222677905 | 0.002297821 |
| TTLL10 | 2.222217319 | 0.005896751 |
| MAFF | 2.22084088 | 4.63E-15 |
| MMRN1 | 2.220169816 | 0.013778629 |
| FHL1 | 2.218065821 | 0.012811521 |
| NFE2L3 | 2.215786889 | 0.00268321 |
| XCL2 | 2.209653366 | 0.000502219 |
| RCAN3 | 2.20798125 | 0.012510035 |
| ENPP1 | 2.193892221 | 0.014171636 |

| | | |
|---|---|---|
| MAN1C1 | 2.190636534 | 0.00312601 |
| IER3 | 2.176477127 | 0.001607825 |
| TTC24 | 2.175570542 | 0.005336525 |
| SCML1 | 2.161265058 | 0.003315952 |
| SPIN3 | 2.149496006 | 0.005300158 |
| PLA2G6 | 2.146855685 | 6.34E-09 |
| TOX2 | 2.138441725 | 0.018405515 |
| LSR | 2.134098322 | 0.007645466 |
| TSPAN4 | 2.109342297 | 0.004707173 |
| BACH2 | 2.102807788 | 1.13E-06 |
| ZMAT4 | 2.095430946 | 0.001208441 |
| PDE4B | 2.090841782 | 6.80E-05 |
| B3GALNT1 | 2.08925033 | 0.015083009 |
| TAF4B | 2.069171566 | 0.017579417 |
| FAM167A | 2.031339399 | 0.002499832 |
| IL18R1 | 2.030167694 | 3.86E-05 |
| SLC17A9 | 2.012849197 | 0.001679691 |
| FUT8 | 2.010414496 | 0.000714077 |
| IFITM3 | 2.005751082 | 9.08E-08 |
| IMMP2L | 2.005050689 | 0.006959766 |
| GNAQ | 1.988506293 | 0.000769341 |
| AXIN2 | 1.985972209 | 0.019014723 |
| FXYD2 | 1.982255418 | 0.004559295 |
| DUSP2 | 1.978657199 | 1.19E-07 |
| SERP2 | 1.978115533 | 0.009072272 |
| FOSB | 1.97565207 | 0.000160107 |
| MYO7A | 1.97543092 | 0.012200394 |
| EPAS1 | 1.965061157 | 0.04575711 |
| LIF | 1.959066583 | 0.010297279 |
| AMPD3 | 1.955846943 | 0.021605623 |
| COL4A4 | 1.951788916 | 0.043535019 |
| SYPL1 | 1.951425481 | 0.010523381 |
| CYGB | 1.943685111 | 0.01524431 |
| SLC44A1 | 1.936263097 | 1.80E-05 |
| LTB | 1.933164129 | 9.86E-05 |
| BEX2 | 1.931430996 | 0.000153228 |
| BCL3 | 1.921142768 | 0.001607825 |
| CHPT1 | 1.908751923 | 0.001105335 |
| TIMP2 | 1.898267337 | 0.013530249 |
| AHR | 1.896876731 | 0.020332567 |
| TIAM1 | 1.888317529 | 0.005018182 |
| CCR7 | 1.876938845 | 0.00826683 |
| TLE3 | 1.868542256 | 0.007054274 |
| SOCS3 | 1.862658014 | 0.037048808 |
| COL24A1 | 1.858366076 | 0.028886317 |

| | | |
|---|---|---|
| ADCY3 | 1.84604287 | 0.032722394 |
| CPNE2 | 1.81502988 | 0.027016474 |
| NFKBIA | 1.802700144 | 4.37E-05 |
| PYROXD2 | 1.762954766 | 0.007702037 |
| GATA3 | 1.705155612 | 2.97E-06 |
| ATP8B4 | 1.700899425 | 0.044069355 |
| GPR183 | 1.69832859 | 0.014089201 |
| UNC93B1 | 1.680391029 | 0.005855446 |
| PLCH2 | 1.67729179 | 0.018405515 |
| CAPN12 | 1.673866558 | 0.003315952 |
| CD55 | 1.627214367 | 0.002821614 |
| CABLES1 | 1.609845504 | 0.005217269 |
| HOXA3 | 1.607788277 | 0.033472557 |
| TC2N | 1.599686252 | 0.007421014 |
| LDLRAP1 | 1.580056387 | 0.014171636 |
| TRAF5 | 1.559587409 | 0.026097189 |
| TMEM123 | 1.545583208 | 0.014999537 |
| BMP2 | 1.544579447 | 0.008685924 |
| PPP1R15A | 1.529215355 | 0.002582424 |
| CLDND1 | 1.501119879 | 0.035904935 |
| GNPTAB | -1.397672739 | 0.022308077 |
| IKZF3 | -1.46092488 | 0.004707173 |
| GZMH | -1.558018265 | 0.030493313 |
| PRR5L | -1.574531396 | 0.013480436 |
| GLIPR1 | -1.595475513 | 0.011761671 |
| FASLG | -1.598250891 | 0.026097189 |
| TMEM2 | -1.609111721 | 0.026097189 |
| LBH | -1.619991604 | 0.000429196 |
| FAM53B | -1.621889172 | 0.000525363 |
| SSBP3 | -1.676350873 | 0.000741457 |
| ARHGEF3 | -1.692061406 | 5.79E-07 |
| COL6A2 | -1.692740822 | 0.018014199 |
| HIPK2 | -1.698068929 | 5.28E-06 |
| MAN1A1 | -1.711469162 | 0.002600225 |
| GRAP2 | -1.721334874 | 0.020250041 |
| TTC38 | -1.726755251 | 0.000773382 |
| NPC1 | -1.75088016 | 0.015363128 |
| LPAR6 | -1.756708615 | 0.007300887 |
| NMUR1 | -1.783636362 | 4.53E-05 |
| RHOBTB3 | -1.785811137 | 0.000664586 |
| TGFA | -1.798867202 | 0.043563421 |
| ABI3 | -1.808950607 | 0.000281162 |
| PLOD1 | -1.809816103 | 0.009800969 |
| NTNG2 | -1.840809686 | 0.033413481 |
| BPGM | -1.846198865 | 0.014171636 |

| | | |
|---|---|---|
| PIF1 | -1.853853395 | 0.022117621 |
| BTBD11 | -1.861729116 | 0.015083009 |
| SLCO4C1 | -1.872590453 | 0.002605464 |
| AK5 | -1.884163401 | 0.029591916 |
| CISH | -1.893913374 | 0.028816662 |
| DRAXIN | -1.894174847 | 0.026858862 |
| PCDH1 | -1.902615233 | 0.0285405 |
| TCF7L2 | -1.90777458 | 0.048488513 |
| CD8B | -1.92457868 | 0.041127969 |
| TBL1X | -1.925432327 | 6.19E-05 |
| ADRB2 | -1.929122656 | 0.007389429 |
| MYRF | -1.933372573 | 0.023775944 |
| PHLDB2 | -1.937553546 | 0.000864418 |
| BOK | -1.975610588 | 0.012171938 |
| CD93 | -2.025433528 | 0.006880661 |
| PRSS57 | -2.026607855 | 0.01429537 |
| EPB41L4A | -2.072942252 | 0.01429537 |
| TNFRSF1A | -2.077318772 | 0.00069523 |
| XPNPEP2 | -2.077478578 | 0.010523381 |
| TSHZ3 | -2.079357982 | 0.010495963 |
| BCL11B | -2.0906216 | 9.14E-12 |
| DGKK | -2.1013329 | 0.000110034 |
| IFI30 | -2.107125196 | 5.21E-06 |
| FCGR3A | -2.148643695 | 4.27E-16 |
| GPR25 | -2.150664786 | 0.007353185 |
| LINGO2 | -2.153314471 | 0.00207457 |
| SYNGR1 | -2.159120744 | 0.000517281 |
| CERCAM | -2.159799991 | 0.006294068 |
| RBPMS2 | -2.171776304 | 0.001982949 |
| ZEB2 | -2.191582818 | 2.13E-08 |
| CCDC65 | -2.19870822 | 0.003233022 |
| PCSK5 | -2.20247926 | 0.000238858 |
| LILRB1 | -2.22674017 | 2.94E-06 |
| MGAM | -2.232446865 | 0.000654202 |
| TNFAIP2 | -2.245728675 | 0.008750116 |
| CAPN5 | -2.248120167 | 0.001515582 |
| ALOX5AP | -2.262801506 | 1.36E-16 |
| FCRL6 | -2.267434399 | 5.50E-11 |
| LYZ | -2.26800786 | 0.001751211 |
| CCL4 | -2.297834076 | 3.90E-19 |
| ERBB2 | -2.306844208 | 7.04E-06 |
| GOLM1 | -2.314485091 | 0.00010819 |
| PRSS23 | -2.320604611 | 7.96E-12 |
| MAF | -2.32833539 | 4.03E-11 |
| ADGRG1 | -2.345740251 | 1.45E-15 |

| | | |
|---|---|---|
| MNDA | -2.346885227 | 0.001412844 |
| ZFYVE28 | -2.351117464 | 9.01E-06 |
| LPAR5 | -2.352527884 | 0.000546423 |
| C16orf45 | -2.384219485 | 0.001005879 |
| KIR2DL3 | -2.384220323 | 1.12E-07 |
| MPEG1 | -2.388057639 | 0.001532392 |
| C10orf128 | -2.390377977 | 0.00014461 |
| PLEKHF1 | -2.418856386 | 1.75E-08 |
| SETBP1 | -2.419281537 | 2.16E-06 |
| TTC16 | -2.433890649 | 0.000153557 |
| KRT72 | -2.44102342 | 7.06E-05 |
| MTSS1 | -2.453761687 | 5.97E-12 |
| GNG2 | -2.463860005 | 2.72E-29 |
| S1PR5 | -2.466360207 | 5.88E-23 |
| KIR3DL1 | -2.485263761 | 1.01E-13 |
| CD1C | -2.490096784 | 0.00108623 |
| GNAL | -2.509943181 | 4.01E-07 |
| ARVCF | -2.51994753 | 8.31E-06 |
| CXCR2 | -2.531096894 | 1.42E-11 |
| RASSF4 | -2.580887986 | 1.91E-11 |
| CCL3 | -2.605064154 | 2.30E-08 |
| ASCL2 | -2.62219014 | 1.86E-07 |
| SYNE2 | -2.628487639 | 4.01E-20 |
| WDFY4 | -2.641463043 | 0.000305324 |
| SLC1A7 | -2.71470013 | 1.27E-05 |
| CXCR1 | -2.73027639 | 3.30E-05 |
| CX3CR1 | -2.753477566 | 4.55E-29 |
| NME8 | -2.777003339 | 1.02E-08 |
| B3GAT1 | -2.779922275 | 8.43E-10 |
| GRK5 | -2.780129439 | 4.24E-09 |
| CTNNA1 | -2.791717998 | 4.71E-07 |
| FGFBP2 | -2.797411579 | 5.22E-23 |
| AGAP1 | -2.807186273 | 4.27E-10 |
| PRDM1 | -2.818351913 | 1.14E-24 |
| LAIR2 | -2.819534636 | 4.51E-10 |
| SPON2 | -2.828259204 | 5.92E-45 |
| PODN | -2.851522976 | 1.01E-05 |
| FGL2 | -2.892758523 | 2.53E-10 |
| CMKLR1 | -2.901480161 | 3.54E-10 |
| FEZ1 | -2.929885497 | 2.51E-10 |
| CACNA2D2 | -2.931520982 | 1.82E-13 |
| AKR1C3 | -2.987859225 | 3.50E-16 |
| KIR2DL1 | -3.076843112 | 6.37E-16 |
| BNC2 | -3.12271759 | 2.03E-18 |
| PTCH1 | -3.161148924 | 1.06E-13 |

| | | |
|---|---|---|
| PALLD | -3.217953235 | 4.09E-16 |
| DAB2 | -3.226720005 | 4.09E-16 |
| KIFC3 | -3.303607692 | 5.36E-14 |
| FCER1A | -3.333784412 | 3.32E-08 |
| CD6 | -3.371135008 | 4.47E-15 |
| LGR6 | -3.547062709 | 2.98E-15 |
| PDGFRB | -3.770734989 | 1.21E-14 |
| PTGDS | -3.805199346 | 1.51E-23 |
| genes with padj < 0.05 and abs(lfcShrink) < 1 | | |
| **DE genes from bulk RNA-Seq ($CD94^{+}CD56^{hi}$ VS $CD94^{+}CD56^{dim}$ )** | | |
| **Gene** | **log2FoldChange** | **padj** |
| IL7R | 3.817231413 | 1.21E-41 |
| FOXC1 | 3.787488805 | 4.98E-08 |
| DUSP4 | 3.461548742 | 1.84E-10 |
| SPINK2 | 3.194542926 | 1.00E-06 |
| COL9A2 | 3.131029701 | 5.94E-09 |
| HAPLN3 | 3.125564446 | 1.02E-06 |
| GNAI1 | 3.053331637 | 4.87E-05 |
| KIT | 2.951081044 | 1.55E-09 |
| NELL2 | 2.940621783 | 2.28E-06 |
| CDHR1 | 2.932659377 | 1.50E-06 |
| CAPG | 2.857379017 | 3.01E-06 |
| SH3BP4 | 2.849002268 | 0.000191263 |
| DACH1 | 2.823707491 | 3.76E-05 |
| NFIX | 2.707412163 | 2.91E-05 |
| GAB1 | 2.650924845 | 4.33E-06 |
| SIRPG | 2.606721567 | 4.87E-05 |
| SPTSSB | 2.591049992 | 1.31E-05 |
| IGFBP4 | 2.585666514 | 0.000154573 |
| PAWR | 2.585341878 | 0.000212197 |
| NR4A3 | 2.570803986 | 5.58E-08 |
| TIMP2 | 2.554316441 | 2.34E-05 |
| ENPP1 | 2.550148292 | 0.000398256 |
| RCAN3 | 2.546414088 | 0.000339777 |
| MMRN1 | 2.53011713 | 0.002664644 |
| TSPAN18 | 2.516205554 | 0.000665263 |
| TNFRSF11A | 2.513906932 | 5.04E-07 |
| PI16 | 2.440992406 | 0.000540832 |
| CDCP1 | 2.422204343 | 0.003306383 |
| SLIT3 | 2.416519667 | 0.003964324 |
| TEC | 2.408020572 | 0.000339777 |
| INPP4B | 2.38838576 | 0.000983333 |
| HOXA9 | 2.380378014 | 0.002227251 |

| | | |
|---|---|---|
| ANO9 | 2.360983198 | 0.00033091 |
| SSBP2 | 2.355742348 | 5.78E-06 |
| LMNA | 2.353468702 | 0.002270555 |
| PPP1R9A | 2.335063925 | 2.85E-07 |
| CCR7 | 2.324837931 | 0.000119075 |
| TTLL10 | 2.323661104 | 0.000495777 |
| KIR2DL4 | 2.319999736 | 0.001143695 |
| FXYD7 | 2.313035368 | 0.000260335 |
| SPRY2 | 2.256846538 | 0.003964937 |
| FHL1 | 2.218790637 | 0.013337631 |
| DENND5A | 2.199798473 | 0.000491467 |
| SCML1 | 2.195931932 | 0.001423147 |
| FLNB | 2.168334763 | 0.000867165 |
| ADGRG3 | 2.15789711 | 0.017411715 |
| TCF7 | 2.109346324 | 1.60E-07 |
| IL12RB2 | 2.082680381 | 2.83E-05 |
| LTB | 2.080709846 | 3.26E-06 |
| RUNX2 | 2.078397253 | 0.00054247 |
| TAF4B | 2.067339575 | 0.022978586 |
| TNFSF11 | 2.057102138 | 0.024412337 |
| PRDM8 | 2.056649239 | 0.006106658 |
| LSR | 2.028218598 | 0.045096718 |
| CRACR2B | 2.022876444 | 0.045839142 |
| DST | 2.021406431 | 0.008689649 |
| C16orf74 | 2.017514714 | 0.014743603 |
| HVCN1 | 2.010643598 | 7.84E-05 |
| ROR1 | 2.007462742 | 0.003363211 |
| GZMK | 1.992181684 | 0.045839142 |
| ZFHX3 | 1.985082363 | 0.004605552 |
| AXIN2 | 1.93917053 | 0.045335739 |
| FXYD2 | 1.935739802 | 0.029271632 |
| TIE1 | 1.935265594 | 0.010338284 |
| DLL1 | 1.924493136 | 0.002899585 |
| COTL1 | 1.921803344 | 0.043847612 |
| SLC44A1 | 1.902395642 | 4.87E-05 |
| CYGB | 1.887187184 | 0.0384865 |
| FOSB | 1.877684003 | 0.000995365 |
| PHOSPHO2 | 1.85113404 | 0.03561554 |
| ZNF521 | 1.851051826 | 0.022141135 |
| SOX4 | 1.841137433 | 0.017508985 |
| INSC | 1.823896588 | 0.025059018 |
| MAFF | 1.799781456 | 1.14E-06 |
| EPHA4 | 1.791531799 | 0.009170543 |
| BACH2 | 1.784012111 | 0.001663516 |
| BEX2 | 1.772440908 | 0.003881153 |

| | | |
|---|---|---|
| CD44 | 1.769279975 | 2.91E-05 |
| FUT8 | 1.759255273 | 0.034717069 |
| NFKBIA | 1.634106832 | 0.00446869 |
| PLA2G6 | 1.611599548 | 0.012727301 |
| IFITM3 | 1.610418169 | 0.009537903 |
| LRP5 | 1.586598719 | 0.049984747 |
| ARHGEF3 | -1.458557709 | 0.006858948 |
| HIPK2 | -1.499615311 | 0.006197222 |
| TMEM171 | -1.500984574 | 0.017411715 |
| LBH | -1.572721829 | 0.002089588 |
| SSBP3 | -1.611536493 | 0.004835745 |
| NMUR1 | -1.653560888 | 0.001907578 |
| GRAP2 | -1.710948198 | 0.031699688 |
| PHLDB2 | -1.723557049 | 0.029016 |
| GZMH | -1.742561252 | 0.000842474 |
| ERBB2 | -1.776545491 | 0.039352887 |
| PLOD1 | -1.821392987 | 0.010066855 |
| MAF | -1.835397477 | 0.000241027 |
| LPAR6 | -1.839294424 | 0.002308701 |
| KRT72 | -1.864013129 | 0.040435053 |
| GOLM1 | -1.86984948 | 0.042779777 |
| BOK | -1.870198159 | 0.038075018 |
| HS6ST1 | -1.891189332 | 0.016302297 |
| FCRL6 | -1.901663709 | 2.06E-05 |
| PODN | -1.921027672 | 0.017411715 |
| RASSF4 | -1.933863059 | 0.000384427 |
| BCL11B | -1.93874756 | 2.18E-08 |
| LINGO2 | -1.954695619 | 0.019146861 |
| TBL1X | -1.979602689 | 2.06E-05 |
| CCDC65 | -1.984919911 | 0.022362634 |
| PRSS23 | -1.987596517 | 2.01E-06 |
| FCGR3A | -1.990957592 | 1.34E-11 |
| LILRB1 | -1.99590055 | 0.000508378 |
| ALOX5AP | -2.003435933 | 3.36E-10 |
| SYNGR1 | -2.0198094 | 0.003881153 |
| TGFA | -2.022901111 | 0.004605552 |
| GPR25 | -2.032213903 | 0.017508985 |
| ZEB2 | -2.046319604 | 2.64E-06 |
| PIF1 | -2.052902853 | 0.005029242 |
| PCDH1 | -2.061988033 | 0.01220424 |
| MTSS1 | -2.090067965 | 1.36E-06 |
| JAKMIP1 | -2.09631697 | 0.004678008 |
| LPAR5 | -2.120794204 | 0.004534276 |
| C10orf128 | -2.125387931 | 0.003557286 |
| CXCR2 | -2.139065573 | 2.28E-06 |

| | | |
|---|---|---|
| CCL4 | -2.170038229 | 2.51E-15 |
| FGL2 | -2.18523166 | 0.000202777 |
| PLEKHF1 | -2.195425495 | 5.31E-06 |
| ADGRG1 | -2.199771349 | 6.62E-12 |
| KIR2DL1 | -2.207515237 | 1.14E-05 |
| SETBP1 | -2.208097026 | 0.000106264 |
| DUSP8 | -2.223942881 | 0.010207089 |
| ZFYVE28 | -2.235471001 | 7.87E-05 |
| LAIR2 | -2.267644238 | 2.52E-05 |
| EPB41L4A | -2.268308467 | 0.002947047 |
| LGR6 | -2.371423511 | 2.06E-05 |
| CCL3 | -2.377201919 | 3.26E-06 |
| ARVCF | -2.385236504 | 6.95E-05 |
| TTC16 | -2.395657659 | 0.00032052 |
| GNG2 | -2.45746383 | 3.28E-29 |
| KIFC3 | -2.479232657 | 2.17E-06 |
| DAB2 | -2.487776787 | 1.39E-07 |
| SPON2 | -2.497041338 | 8.04E-30 |
| FGFBP2 | -2.513825459 | 4.91E-16 |
| S1PR5 | -2.514685615 | 1.99E-24 |
| CX3CR1 | -2.529312104 | 6.63E-22 |
| B3GAT1 | -2.548117138 | 3.28E-07 |
| NME8 | -2.557318827 | 8.80E-07 |
| AGAP1 | -2.56055033 | 1.70E-07 |
| SYNE2 | -2.571384105 | 1.43E-18 |
| CD6 | -2.590688978 | 2.44E-07 |
| PRDM1 | -2.599720153 | 5.18E-19 |
| BNC2 | -2.626920383 | 5.74E-11 |
| CMKLR1 | -2.645501813 | 1.38E-07 |
| CXCR1 | -2.650838651 | 7.72E-05 |
| CTNNA1 | -2.652180766 | 4.21E-06 |
| FEZ1 | -2.666536761 | 1.19E-07 |
| AKR1C3 | -2.666570923 | 2.70E-11 |
| PALLD | -2.681178474 | 1.55E-09 |
| ASCL2 | -2.705445337 | 5.58E-08 |
| PTGDS | -2.739711248 | 1.25E-09 |
| GRK5 | -2.803256643 | 4.12E-09 |
| CACNA2D2 | -2.81713462 | 9.96E-12 |
| PTCH1 | -3.050534628 | 4.50E-12 |
| PDGFRB | -3.142471273 | 1.55E-09 |
| genes with padj < 0.05 and abs(lfcShrink) < 1 | | |
| **DE genes from bulk RNA-Seq ($CD94^{+}CD56^{dim}$ VS $CD94^{-}CD56^{dim}$ )** | | |
| **Gene** | **log2FoldChange** | **padj** |

| | | |
|---|---|---|
| KLRC1 | 2.81082892 | 9.05E-08 |
| KLRC2 | 2.413131496 | 7.29E-05 |
| CD1C | -2.45437148 | 0.030172661 |
| FCER1A | -3.091245955 | 0.000250858 |
| genes with padj < 0.05 and abs(lfcShrink) < 1 | | |
